# BRI2-mediated regulation of TREM2 processing in microglia and its potential implications for Alzheimer’s disease and related dementias

**DOI:** 10.1101/2023.06.14.544924

**Authors:** Tao Yin, Luciano D’Adamio

## Abstract

*ITM2B/BRI2* mutations cause familial forms of Alzheimer’s disease (AD)-related dementias by disrupting BRI2’s protein function and leading to the accumulation of amyloidogenic peptides. Although typically studied in neurons, our findings show that BRI2 is highly expressed in microglia, which are crucial in AD pathogenesis due to the association of variants in the microglial gene TREM2 with increased AD risk. Our single-cell RNAseq (scRNAseq) analysis revealed a microglia cluster that depends on a Trem2 activity that is inhibited by Bri2, pointing to a functional interaction between *Itm2b/Bri2* and *Trem2*. Given that the AD-related Amyloid-β Precursor protein (APP) and TREM2 undergo similar proteolytic processing, and that BRI2 inhibits APP processing, we hypothesized that BRI2 may also regulate TREM2 processing. We found that BRI2 interacts with Trem2 and inhibits its processing by α-secretase in transfected cells. In mice lacking Bri2 expression, we observed increased central nervous system (CNS) levels of Trem2-CTF and sTrem2, which are the products of α-secretase processing of Trem2, indicating increased Trem2 processing by α-secretase *in vivo*. Reducing Bri2 expression only in microglia resulted in increased sTrem2 levels, suggesting a cell-autonomous effect of Bri2 on α-secretase processing of Trem2. Our study reveals a previously unknow role of BRI2 in regulating TREM2-related neurodegenerative mechanisms. The ability of BRI2 to regulate the processing of both APP and TREM2, combined with its cell-autonomous role in neurons and microglia, makes it a promising candidate for the development of AD and AD-related dementias therapeutics.

## Introduction

*ITM2B* mutations have been linked to four autosomal dominant neurodegenerative diseases, the Familial British Dementia (FBD)^1^, the Familial Danish Dementia (FDD)^2^ and the newly discovered Familial Chinese^3^ and Familial Korean Dementias^4^. *ITM2B* encodes for a type II membrane protein called BRI2. BRI2 is synthesized as a precursor protein that is cleaved at the C-terminus by proprotein convertase into mature BRI2 and a 23 amino acid-long (Bri23) soluble C-terminal fragment^5^. All pathogenic *ITM2B* mutations lead to changes in the C-terminal region of BRI2, resulting in the production of longer C-terminal fragments, which are processed into amyloidogenic peptides.

FDD and FBD share similarities with AD in terms of their histopathological features, such as, neuroinflammation, neurodegeneration, the presence of extracellular amyloid plaques and intraneuronal neurofibrillary tangles. However, the composition of the amyloid plaques in FDD and FBD is different from that of AD. In AD, the amyloid plaques are primarily composed of amyloid-β (Aβ) peptides, which derive from the proteolytic processing of APP, whereas in FDD and FBD, the plaques are composed of the cleavage products of the mutant BRI2 proteins, the ADan peptide, or the ABri peptide, respectively^6^. Of note, in patients with FDD, Aβ deposition was observed either in combination with ADan or alone^2^. These differences in the composition of the plaques have led to the classification of FDD and FBD as distinct neurodegenerative diseases and to the conclusion that *ITM2B* mutations cause accumulation of amyloidogenic peptides aggregates, which lead to neuronal damage and dementia.

Findings from studies on *Itm2b-KO* and conditional *Itm2b-KO* mice, have shown that BRI2 has a cell autonomous physiological function in synaptic transmission and plasticity in glutamatergic neurons at both presynaptic and postsynaptic termini^7^. FDD and FBD knock-in rodents show synaptic plasticity deficits like those observed in *Itm2b-KO* mice^8, 9^. In addition, in FDD and FBD knock-in animal models, the mutant forms of BRI2 protein have been found to be unstable and rapidly degraded^8, 10, 11^. These findings suggest that the pathogenesis of FDD and FBD may be more complex than originally thought and that both the accumulation of amyloidogenic peptides and a loss of BRI2 protein function may contribute to the development of these diseases.

BRI2 has also a dual anti-amyloidogenic function, reducing both Aβ production and Aβ aggregation. It has been found that BRI2 binds to APP in cis, thereby reducing APP cleavage and Aβ production^12–15^. Additionally, the extracellular domain of BRI2 includes a BRICHOS domain that inhibits or delays Aβ aggregation^16, 17^. These activities of BRI2 on APP processing and Aβ aggregation are supported by evidence that APP and APP processing play a role in long-term synaptic plasticity and memory deficits in FDD and FBD knock-in mice^15, 18–21^.

While BRI2 function has been primarily studied in neuronal cells, BRI2 may also have biological roles also in other CNS cell types. Analysis of mouse and human nervous system scRNAseq^22^ and single nuclei RNAseq^23^ (snRNAseq) data showed that *ITM2B* expression in the CNS is highest in microglia (Figure 1). This finding is significant as increasing evidence link neuroinflammation to AD^24, 25^. For instance, variants of the TREM2 gene, which is exclusively expressed in microglia^26^, have been shown to increase the risk of developing sporadic, late-onset AD^27^. TREM2 also undergoes regulated intramembrane proteolysis^28^, similar to APP, in which α-secretase cleaves TREM2^29^ resulting in the release of soluble TREM2 ectodomain (sTREM2) and the membrane-tethered C-terminal fragment (TREM2-CTF). Levels of sTREM2 are increased in the CSF and CNS soluble fraction of AD patients^30–32^, suggesting a potential role of TREM2 processing by α-secretase in AD pathology. TREM2-CTF is subsequently cleaved in the transmembrane region by γ-secretase^28^.

**Figure 1.**
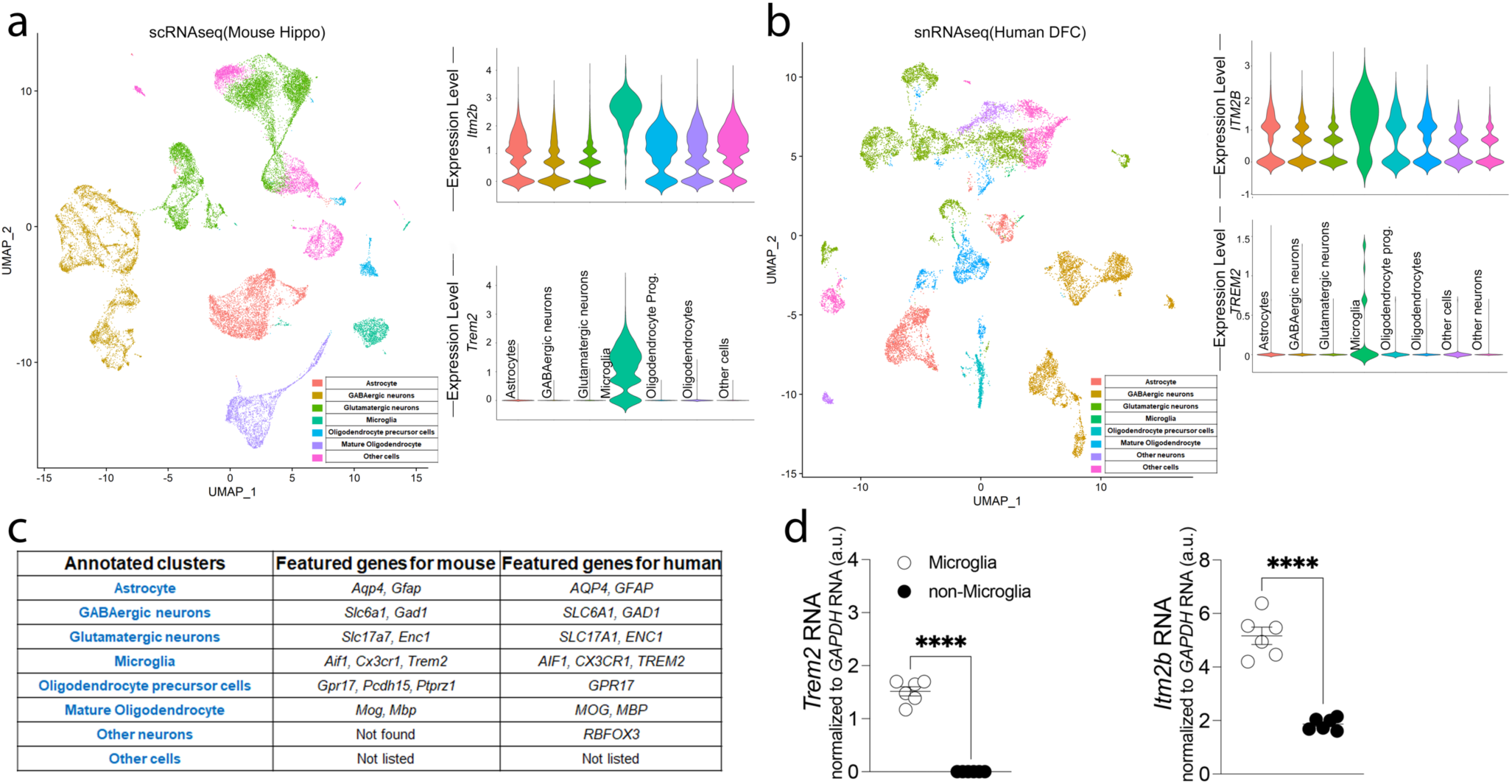
Analysis of *Itm2b* mRNA expression in the CNS. UMAP visualization of mouse hippocampal cell clusters (a, left panel) and human DFC cell clusters (b, left panel), classified by cell type based on DEG identified by Seurat v4. (b) Violin plots represent the log-normalized expression of *Itm2b* and *Trem2* across cell populations in mouse hippocampal cell clusters (a, right panel) and human DFC cell clusters (b, right panel). (c) *Itm2b* and *Trem2* mRNA expression in mouse microglia and non-microglia cells analyzed by quantitative RT-PCR.

Based on the above evidence, in the present study, we investigated the role of BRI2 in microglia, with a focus on potential BRI2-TREM2 physiological interactions mirroring those observed between BRI2 and the other AD-related secretases’ substrate, APP. Understanding the functions of BRI2 in microglia and its interactions with other AD-related proteins may provide further insight into the complex pathogenic mechanisms underlying AD and related dementias.

## Results

### In the CNS, microglia express the highest levels of *Itm2b* mRNA

Unbiased clustering with high resolution (1.0) of mouse scRNAseq^22^ and human snRNAseq^23^ data sets revealed 40 clusters in the mouse hippocampus and 31 clusters in the human dorso-lateral prefrontal cortex (DFC). Similar clusters were then manually grouped together when visualizing with uniform manifold approximation and projection (UMAP)^33^ to simplify the cell-types annotation (Figure 1 a, left panel, and 1b, left panel). Specifically, we combined clusters corresponding to (1) Astrocytes; (2) GABAergic neurons; (3) glutamatergic neurons, (4) microglia, (5) oligodendrocyte precursor cells, (6) mature oligodendrocytes and (7) other nonspecific neurons or (8) other cells. The major populations were annotated based on differential expression of known cell-type-specific marker genes (listed in Figure 1c). *Itm2b* and *Trem2* mean mRNA expression levels were greater in the cluster identified as microglia compared to the rest of the cells in this both mouse and human datasets (Figure 1a, right panel, and 1b, right panel).

To validate these findings, CD11b^+^ cells were isolated employing the microglia isolation protocol^34^ using the Adult Brain Dissociation Kit and the CD11b magnetic microbeads from Miltenyi. Prior to brain harvesting, we removed peripheral myeloid cells and blood from brain tissue via intracardiac catheterization and perfusion. Quantitative RT-PCR analysis showed that the microglia-specific marker *Trem2* mRNA was expressed in the purified microglia but not in the unbound flow-through cells (referred to as non-microglia), indicating the purity and efficiency of the microglia isolation, and that *Itm2b* mRNA levels are significantly higher in microglia than non-microglia (Figure 1d). Taken together, these findings indicate that *Itm2b* expression in the CNS is predominantly in microglia.

### *Itm2b* modulates microglial transcriptome in a *Trem2*-dependent manner

Next, we examined the impact of *Itm2b*, *Trem2* and combined *Itm2b*-*Trem2* deficiency on microglial transcriptome by scRNAseq. Single/live CD11b^+^ cells were isolated from WT control, *Itm2b-KO*, *Trem2-KO* and *Itm2b/Trem2-dKO* (double KO) brains, and single-cell transcriptomes were generated using the 10x Genomics platform in two independent experiments. After quality control, cells were plotted on UMAP dimensions for visualization (Supplementary Figure 1a, left panels). To specifically select microglia for further analysis, we performed cell type annotation using a single cell dataset published by Van Hove^35^ as a reference. Cells predicted to be of the type “microglia” with > 95% confidence were retained for further analysis (Supplementary Figure 1b). As the samples were sequenced in two different experiments (Data 1 and Data 2), the scRNA dataset integration functionality of the Seurat package was used to perform the joint analysis. Select sample datasets indicated above from Data 1 and Data 2 were integrated using the first 20 principal components into Object1 containing information on 297,215 cells (Supplementary Figure 1c). Unsupervised clustering of microglia revealed a total of 16 distinct microglia clusters across all mice (Figure 2a, Supplementary Figure 1c, Supplementary Figure 2a and 2b). Based on expression of specific marker genes^36^, we identified cluster 6 as MHC-II microglia (high expression of genes such as H2-Aa, H2-Ab1, H2-DMb1, H2-DMb2, H2-DMa and Cd74, Figure 2b). Cluster 10 was identified as IFN-R microglia based on the high expression of genes such as Ifitm3, Isg15 and Ifit3 (Figure 2b). Clusters 12 was characterized by high expression of microglia activation genes *Apoe* and *Lyz2* and is referred to as Activated cluster. The gene expression heatmap showing the top five enriched genes for each microglia cluster and the number of cells per cluster is shown in Supplementary Figure 2c.

**Figure 2.**
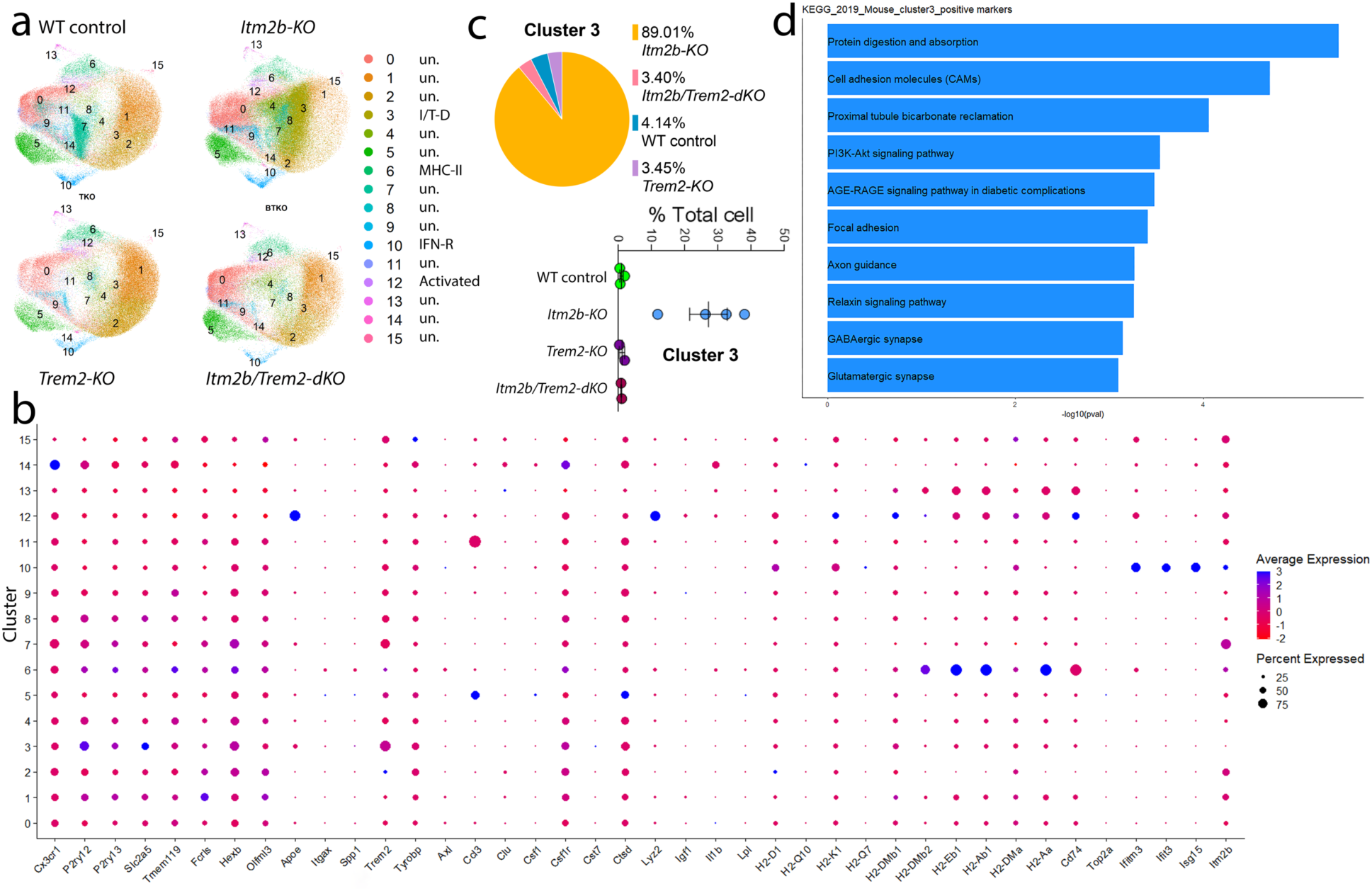
*Itm2b* modulates microglial transcriptome and clusterization in a *Trem2*-dependent manner. (a) UMAPs of microglia grouped by genotype. (b) Average scaled expression levels of selected signature genes per cluster and cluster’s annotation based on expression of signature genes. (c) Proportional contribution of each genotype and proportional contribution of individual samples of each genotype to cluster 3. (d) KEGG pathway enrichment analysis of pathways upregulated in cluster 3.

We next examined the representation of these microglia clusters in each genotype (Supplementary Figure 2d). Although several clusters showed a differential representation in different genotypes, cluster 3 displayed the most distinctive pattern, with a nearly exclusive representation in *Itm2b-KO* mice, as 89% of the microglia in this cluster were derived from *Itm2b-KO* mice (Figure 2c and Supplementary Figure 2d). In contrast, only a minor contribution was observed from other genotypes, including WT animals, *Trem2-KO*, and *Itm2b/Trem2-dKO* mice. *Itm2b* was one of the most differentially expressed genes in cluster 3, with a significant downregulation compared to other clusters (Supplementary Figure 2c and Figure 2b), consistent with the evidence that *Itm2-KO* cells were highly represented in this cluster. Analysis of the relative abundance of cluster 3 cells in each single animal revealed that in all 4 *Itm2b-KO* mice, microglia assigned to cluster 3 were abundant, ranging from 12.1% to 38.3% of total microglia (Figure 2c). In contrast, the percentage of microglia assigned to this cluster in WT animals, *Trem2-KO*, and *Itm2b/Trem2-dKO* mice was minimal, ranging from 0.5% to 1.2% (Figure 2c). KEGG pathway enrichment analysis indicated that several neuronal function-related pathways were up-regulated in cluster 3 relative to all other clusters. These include pathways related to axon guidance, GABAergic and Glutamatergic synapses (Figure 2d and Table 1). Overall, these data demonstrate that the observed effect is not related to animal-specific factors, but rather to genotype-specific factors.

**Table 1.**
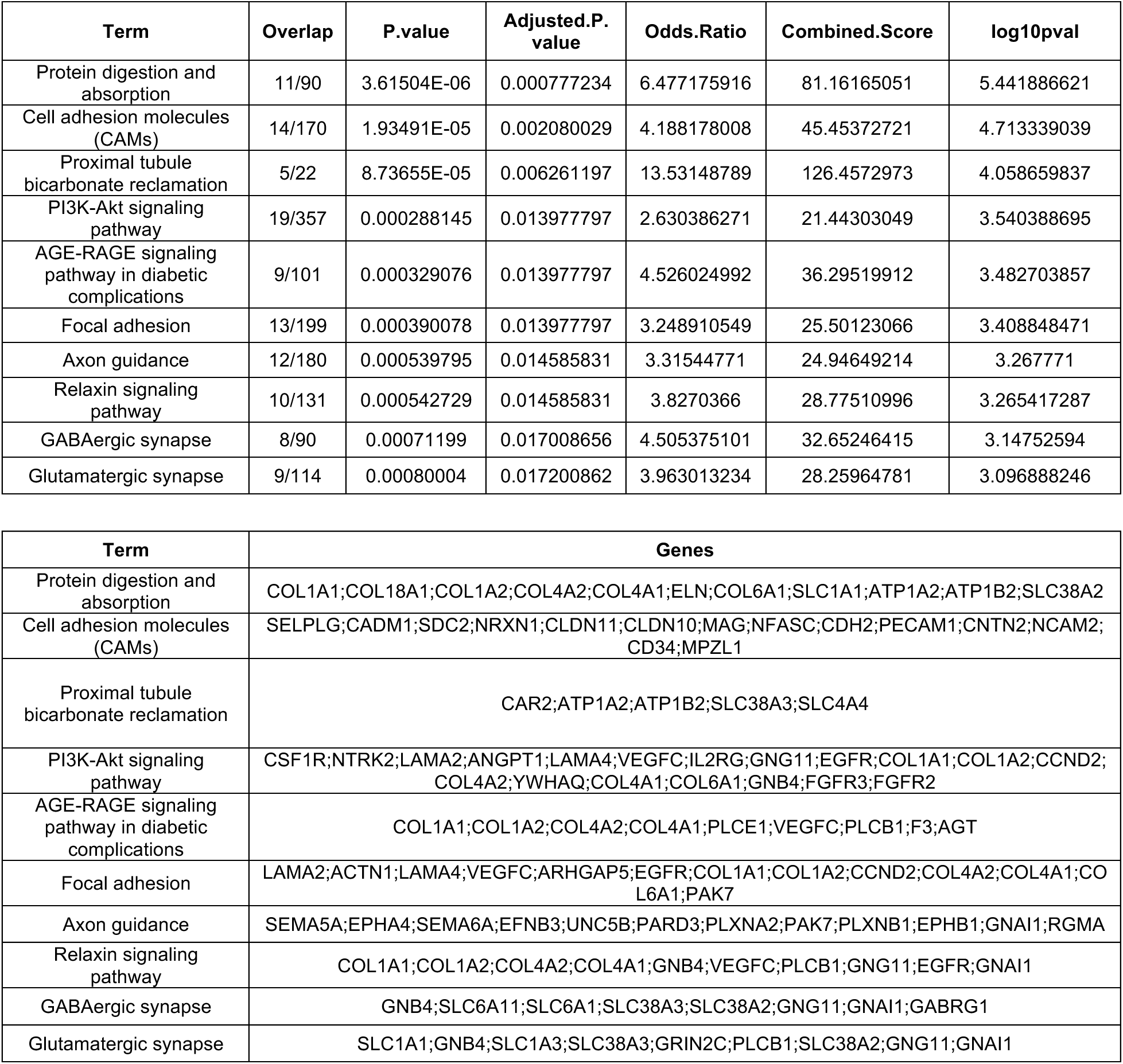
KEGG pathway enrichment analysis indicated that several neuronal function-related pathways were up-regulated in cluster 3 relative to all other clusters.

| Term | Overlap | P.value | Adjusted.P.<br>value | Odds.Ratio | Combined.Score | log10pval |
| --- | --- | --- | --- | --- | --- | --- |
| Protein digestion and absorption | 11/90 | 3.61504E-06 | 0.000777234 | 6.477175916 | 81.16165051 | 5.441886621 |
| Cell adhesion molecules (CAMs) | 14/170 | 1.93491E-05 | 0.002080029 | 4.188178008 | 45.45372721 | 4.713339039 |
| Proximal tubule bicarbonate reclamation | 5/22 | 8.73655E-05 | 0.006261197 | 13.53148789 | 126.4572973 | 4.058659837 |
| PI3K-Akt signaling pathway | 19/357 | 0.000288145 | 0.013977797 | 2.630386271 | 21.44303049 | 3.540388695 |
| AGE-RAGE signaling pathway in diabetic complications | 9/101 | 0.000329076 | 0.013977797 | 4.526024992 | 36.29519912 | 3.482703857 |
| Focal adhesion | 13/199 | 0.000390078 | 0.013977797 | 3.248910549 | 25.50123066 | 3.408848471 |
| Axon guidance | 12/180 | 0.000539795 | 0.014585831 | 3.31544771 | 24.94649214 | 3.267771 |
| Relaxin signaling pathway | 10/131 | 0.000542729 | 0.014585831 | 3.8270366 | 28.77510996 | 3.265417287 |
| GABAergic synapse | 8/90 | 0.00071199 | 0.017008656 | 4.505375101 | 32.65246415 | 3.14752594 |
| Glutamatergic synapse | 9/114 | 0.00080004 | 0.017200862 | 3.963013234 | 28.25964781 | 3.096888246 |

| Term | Genes |
| --- | --- |
| Protein digestion and absorption | COL1A1;COL18A1;COL1A2;COL4A2;COL4A1;ELN;COL6A1;SLC1A1;ATP1A2;ATP1B2;SLC38A2 |
| Cell adhesion molecules (CAMs) | SELPLG;CADM1;SDC2;NRXN1;CLDN11;CLDN10;MAG;NFASC;CDH2;PECAM1;CNTN2;NCAM2;CD34;MPZL1 |
| Proximal tubule bicarbonate reclamation | CAR2;ATP1A2;ATP1B2;SLC38A3;SLC4A4 |
| PI3K-Akt signaling pathway | CSF1R;NTRK2;LAMA2;ANGPT1;LAMA4;VEGFC;IL2RG;GNG11;EGFR;COL1A1;COL1A2;CCND2;COL4A2;YWHAQ;COL4A1;COL6A1;GNB4;FGFR3;FGFR2 |
| AGE-RAGE signaling pathway in diabetic complications | COL1A1;COL1A2;COL4A2;COL4A1;PLCE1;VEGFC;PLCB1;F3;AGT |
| Focal adhesion | LAMA2;ACTN1;LAMA4;VEGFC;ARHGAP5;EGFR;COL1A1;COL1A2;CCND2;COL4A2;COL4A1;COL6A1;PAK7 |
| Axon guidance | SEMA5A;EPHA4;SEMA6A;EFNB3;UNC5B;PARD3;PLXNA2;PAK7;PLXNB1;EPHB1;GNAI1;RGMA |
| Relaxin signaling pathway | COL1A1;COL1A2;COL4A2;COL4A1;GNB4;VEGFC;PLCB1;GNG11;EGFR;GNAI1 |
| GABAergic synapse | GNB4;SLC6A11;SLC6A1;SLC38A3;SLC38A2;GNG11;GNAI1;GABRG1 |
| Glutamatergic synapse | SLC1A1;GNB4;SLC1A3;SLC38A3;GRIN2C;PLCB1;SLC38A2;GNG11;GNAI1 |

The enzymatic dissociation protocol employed to isolate microglia from brains has the potential to activate microglia, and thus, the scRNAseq data may not fully represent the actual microglia populations in the brains of the mutant mice analyzed. Nevertheless, our findings strongly support the existence of a functional interaction between *Itm2b* and *Trem2*, with *Itm2b* potentially acting epistatically to *Trem2* to regulate cluster 3 expression. This could be due to the inhibition of a Trem2 function by Bri2, which is relieved in the absence of Bri2, leading to the expansion of clusters 3. Deletion of Trem2 or both Trem2 and Bri2 does not result in an increase in clusters 3, indicating Trem2’s essential role in this pathway and Bri2’s role as an inhibitory regulator. Thus, we have named this cluster as the *Itm2b*-*Trem2* dependent cluster (I/T-D).

### BRI2 and TREM2 interact in transiently transfected cells

Like APP, TREM2 is processed by the α- and the γ-secretases. Although the functional consequences of TREM2 processing are not well understood, TREM2 cleavage has been suggested to play a role in regulating the activity of microglia in the brain as well as in AD pathogenesis^30–32, 37^. BRI2 interacts with APP via its membrane-proximal region, which contains the secretases’ cleavage sites, and inhibits APP processing^14, 20^. If APP and TREM2 share structural similarities in these regions, it is possible that BRI2 also interacts with TREM2 and inhibit its processing in a similar manner.

To test these hypotheses, we transfected HEK293 and N2a cells with constructs coding for BRI2 FLAG-tagged at the NH_2_-terminal cytoplasmic tail (F-BRI2) and rat Trem2 (Trem2-Miα isoform, UniProtKB - A0A6G8MV71)^38^, and then immunoprecipitated the lysates with an anti-FLAG antibody to pull down BRI2. The immunoprecipitants were analyzed using a Trem2-specific antibody to detect any interaction between BRI2 and Trem2. The results of the experiment showed that Trem2 was precipitated by the anti-FLAG antibody only when BRI2 and Trem2 were co-expressed (Figure 3a). This suggests a direct interaction between BRI2 and Trem2.

**Figure 3.**
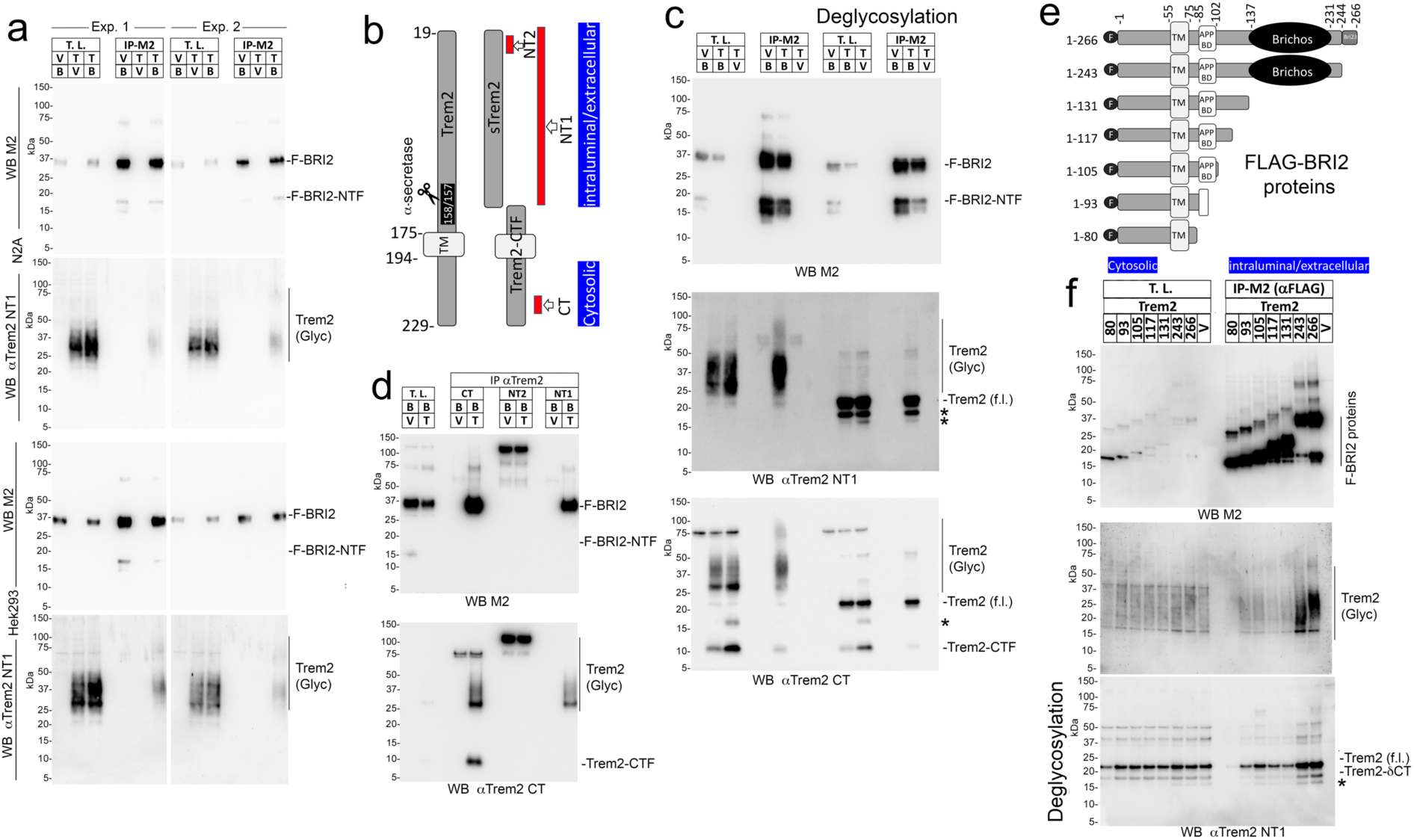
BRI2 binds Trem2 in transfected cells. (a) N2A or HEK293 cells were transfected with F-BRI2 (B) and Trem2 (T), either alone or in combination (V=empty pcDNA3.1vector) and analyzed by Western blot with anti-FLAG (M2) and anti-Trem2 (NT1) on total lysates (T.L.) and M2 immunoprecipitants (IP-M2). For each cell line, two independent transfections were performed (Exp. 1 and Exp. 2). (b) Schematic representation of Trem2 and the two products of α-secretase cleavage, sTrem2, and Trem2-CTF. TM indicates the transmembrane region of Trem2. Red bars point to the antigenic regions used to produce the anti-Trem2 antibodies CT, NT1 and NT2. The cytosolic and intralumenal/extracellular regions of Trem2 are indicated. (c) Western blot analysis with anti-FLAG, anti-Trem2 NT1, and anti-Trem2 CT antibodies of T.L. and IP-M2 from HEK293 cells transfected with F-BRI2 and Trem2, either alone or in combination, with or without deglycosylation. (d) Western blot analysis with anti-FLAG and anti-Trem2 CT antibodies of immunoprecipitants obtained with CT, NT1, and NT2 antibodies from HEK293 cells expressing either F-BRI2 alone or F-BRI2 plus Trem2. (e) Schematic representation of the F-BRI2 constructs used in (f). The Bri23 region, transmembrane region (TM), Brichos domain, APP-binding domain (APP BD), FLAG tag (F), cytosolic and intralumenal/extracellular regions are indicated. (f) WB analysis with anti-FLAG and anti-Trem2 antibodies of lysates and immunoprecipitants from HEK293 cells expressing F-BRI2 deletion mutants plus Trem2 or Trem2 alone (V). *Indicates Trem2 species of unclear primary structure.

Trem2 is highly glycosylated, resulting in heterogeneous sizes (Figure 3a and 3c). Deglycosylation of Trem2 leads to a homogeneous protein of about 22 kDa, which is efficiently immunoprecipitated by the anti-FLAG antibody in cells co-expressing F-BRI2 and Trem2 (Figure 3c, middle panel). The anti-Trem2 NT1 antibody (the NT1 epitope is depicted in Figure 3b) used in the study does not recognize Trem2-CTF, the membrane-bound product of Trem2 processing by α-secretase. However, the anti-Trem2 CT antibody (the epitope of CT is depicted in Figure 3b), detected both Trem2 and Trem2-CTF in both total lysates and immunoprecipitants (Figure 3c), indicating that F-BRI2 interacts with both Trem2 and Trem2-CTF.

Next, we performed a reverse immunoprecipitation by using antibodies against Trem2 to pull down BRI2. We found that both anti-Trem2 CT and anti-Trem2 NT1 antibodies were able to pull down F-BRI2 only when Trem2 was co-expressed with BRI2 (Figure 3d). However, a different antibody (anti-Trem2 NT2, see epitope in Figure 3b) that did not immunoprecipitated Trem2 was not able to pull down BRI2 (Figure 3d), indicating that the interaction between BRI2 and Trem2 is specific. This confirms that BRI2 and Trem2 interact with each other, suggesting a functional interaction between the two proteins.

To define the domain(s) of BRI2 that bind(s) to Trem2, BRI2 fragments progressively deleted from the COOH-terminus (scheme in Figure 3e) were co-transfected with Trem2 in HEK293 cells. Trem2 was expressed at similar levels in all transfections (Figure 3f, upper panel). Deletion of the BRI2-BRICHOS domain (F-BRI2_1-131_) reduced binding of Trem2. F-BRI2_1-117_, F-BRI2_1-105_, and F-BRI2_1-93_ bound Trem2 with similar efficiency to F-BRI2_1-131_, while F-BRI2_1-80_ did not bind to Trem2 (Figure 3f). This suggests the presence of two Trem2-binding domains in BRI2; one probably contained in the BRI2-BRICHOS domain and the other between amino acids 81 and 93 of BRI2. This second domain partially overlaps with the APP-binding domain of BRI2 (Figure 3e).

### BRI2 reduces α-secretase-mediated processing of TREM2 in transiently transfected cells

Binding of BRI2 to APP has been shown to reduce secretases-mediated processing of APP^14^. To test if BRI2 has a similar effect on Trem2 processing, HEK293 cells were co-transfected with Trem2 and either empty vector or F-BRI2. Co-transfection of Trem2 with F-BRI2 in HEK293 cells led to an increase in Trem2 levels and a decrease in Trem2-CTF in the cell lysates and sTrem2 in the tissue culture media (Figure 4a and 4b). The observation that overexpression of BRI2 leads to elevated levels of the α-secretase substrates Trem2, and a simultaneous decrease in the α-secretase products Trem2-CTF and sTrem2, strongly supports the idea that BRI2 functions as an inhibitor of Trem2 processing by α-secretase.

**Figure 4.**
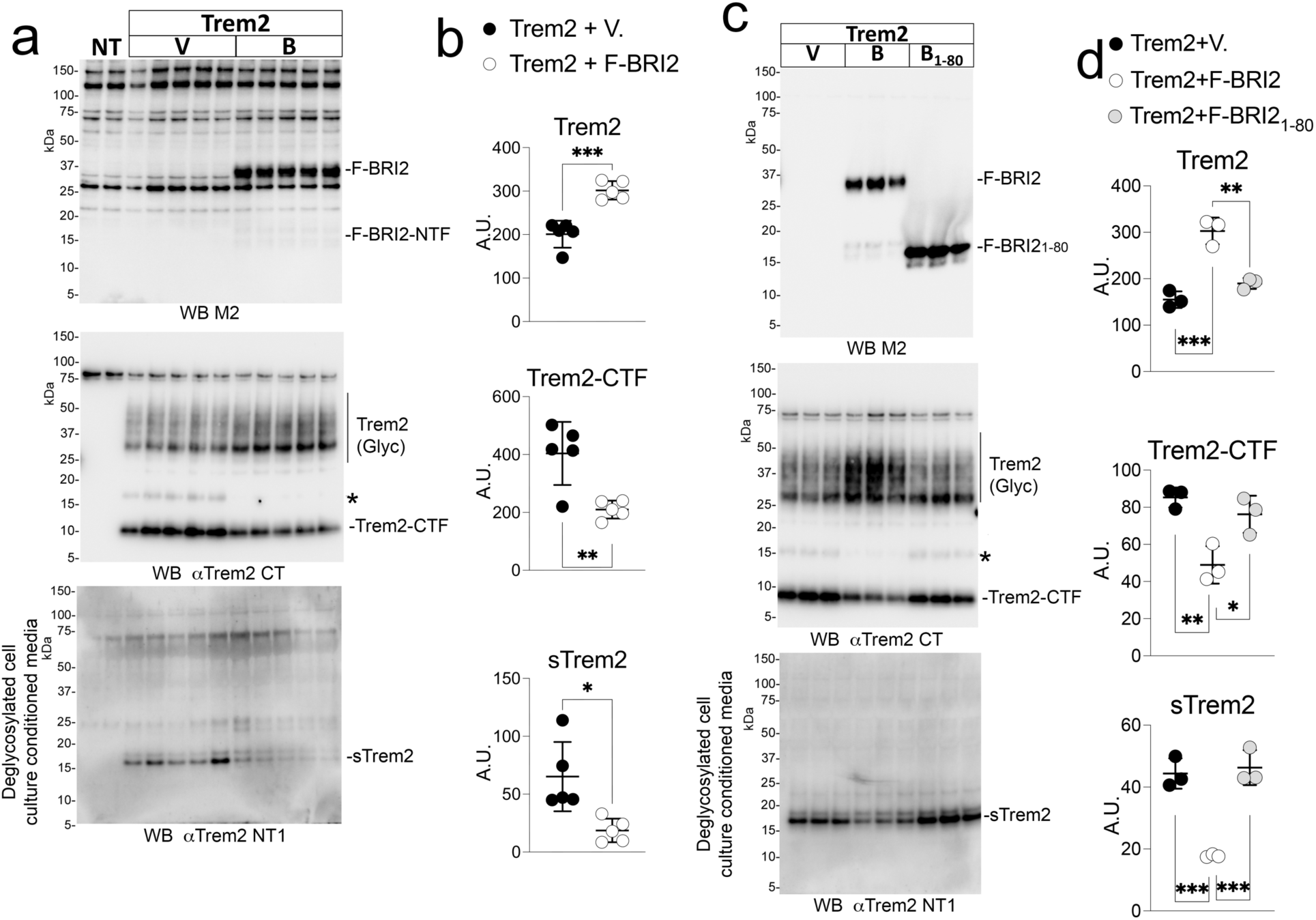
BRI2 reduces α-secretase cleavage of Trem2 in transfected cells. (a) HEK293 cells were transfected with Trem2 and either empty vector (V) or F-BRI2 (B). NT are non-transfected cells. Western blot of cell lysates with either the anti-FLAG antibody M2 or the anti-Trem2 antibody CT. Western blot of deglycosylated culture supernatants with the anti-Trem2 antibody NT1. (b) Quantification of Trem2, Trem2-CTF and sTrem2 levels detected by Western blot in (a). Data were analyzed by two-tailed unpaired t test: Trem2, P=0.0003; Trem2-CTF, P=0.005; sTrem2. (c) HEK293 cells were transfected with Trem2 and either empty vector (V), F-BRI2 or deletion mutant F-BRI2_1-80_. Western blot of cell lysates with either the anti-FLAG antibody M2 or the anti-Trem2 antibody CT. Western blot of deglycosylated culture supernatants with the anti-Trem2 antibody NT1 (lower panel). (d) Quantification of Trem2, Trem2-CTF and sTrem2 levels detected by Western blot in (c). Data were analyzed by ordinary one-way ANOVA followed by post-hoc Tukey’s multiple comparisons test when ANOVA showed significant differences. Trem2 F (2, 6) = 41.86, P=0.0003; post-hoc Tukey’s multiple comparisons test: Trem2+V vs. Trem2+F-BRI2, P=0.0003; Trem2+V vs. Trem2+F-BRI2_1-80_, P=0.1786; Trem2+F-BRI2 vs. Trem2+F-BRI2_1-80_, P=0.0013. Trem2-CTF F (2, 6) = 13.92, P=0.0056; post-hoc Tukey’s multiple comparisons test: Trem2+V vs. Trem2+F-BRI2, P=0.0055; Trem2+V vs. Trem2+F-BRI2_1-80_, P=0.4600; Trem2+F-BRI2 vs. Trem2+F-BRI2_1-80_, P=0.0210. sTrem2 F (2, 6) = 40.57, P=0.0003; post-hoc Tukey’s multiple comparisons test: Trem2+V vs. Trem2+F-BRI2, P=0.0007; Trem2+V vs. Trem2+F-BRI2_1-80_, P=0.8574; Trem2+F-BRI2 vs. Trem2+F-BRI2_1-80_, P=0.0005. All data are shown as means +/-SEM: *=P<0.05, **=P<0.01, ***=P<0.001.

To determine whether BRI2 binding is required for the effects on Trem2 processing, HEK293 cells were co-transfected with Trem2 and either empty vector, F-BRI2, or F-BRI2_1-80_ that does not bind Trem2 (Figure 3f). WB analysis shows that F-BRI2 significantly increased Trem2 levels, reducing Trem2-CTF and sTrem2 amounts (Figure 4c and 4d). In contrast, F-BRI2_1-80_ did not alter levels of Trem2, Trem2-CTF and sTrem2 (Figure 4c and 4d). This implies that BRI2’s inhibition of α-secretase processing of Trem2 requires its binding to Trem2.

### *Itm2b* deletion results in elevated CNS levels of Trem2-CTF and sTrem2 in mice

To confirm the *in vitro* findings, we measured the levels of Trem2, sTrem2, and Trem2-CTF in the central nervous system of approximately 245-day-old *Itm2b-KO* and WT mice. Due to the heterogeneity of Trem2 and sTrem2 caused by glycosylation, and their limited expression in microglia (which represent only approximately 10% of CNS cells), we employed two ELISAs: ELISA 1, which detects only Trem2, and ELISA 2, which detects both Trem2 and sTrem2, to accurately measure their levels in vivo (Figure 5a). Brain homogenates were separated into two fractions using centrifugation at 100000g. The pellet fraction (P100) is enriched in cells-derived material, while the soluble fraction (S100) is enriched in soluble extracellular material. Using ELISA 1 (Figure 5b, upper panels) we found that Trem2 is not detectable in *Trem2-KO* brains, demonstrating the specificity of the assay. Furthermore, Trem2 was detected only in the P100 fraction, indicating that it is cell-bound. We did not observe significant differences in the levels of Trem2 between *Itm2b-KO* and control WT animals. ELISA 2 (Figure 5b, lower panels) revealed that Trem2 proteins are not detectable in *Trem2-KO* brains, confirming the specificity of the assay. We found that sTrem2 was significantly enriched in the S100 fraction of *Itm2b-KO* animals compared to control WT animals.

**Figure 5.**
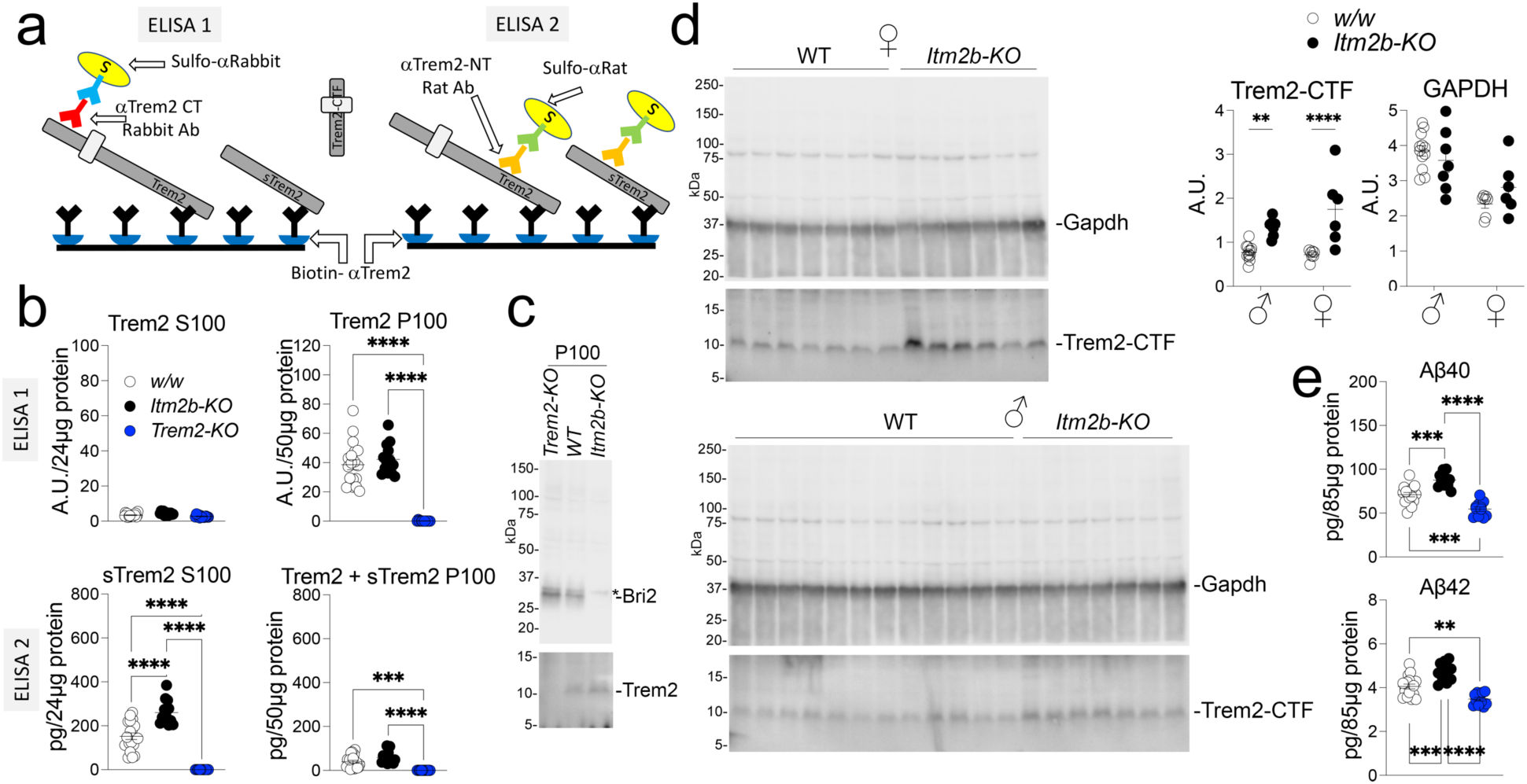
Loss of Bri2 leads to increased CNS levels of sTrem2 and Trem2-CTF. (a) Schematic representation of ELISA 1 and ELISA 2. Both ELISAs use the same Biotinylated-αTrem2 capture antibody (in black). ELISA 1 uses αTrem2-CT (red) + Sulfo-αRabbit (blue) detection antibodies. ELISA 2 uses αTrem2-NT (orange) + Sulfo-αRat (green) detection antibodies. Trem2 can be detected by both ELISAs, sTrem2 can be detected only by ELISA 2: neither ELISA can detect Trem2-CTF. (b) Quantification of Trem2 and sTrem2 in the P100 and S100 brain fractions of ∼245 days old control (*w/w*, 7 females and 12 males), *Itm2b-KO* (6 females and 7 males) and *Trem2-KO* (6 females and 7 males) mice. Data were analyzed by ordinary one-way ANOVA followed by post-hoc Tukey’s multiple comparisons test when ANOVA showed significant differences. ELISA1, Trem2 in P100 fraction F (2, 42) = 60.10, P<0.0001; post-hoc Tukey’s multiple comparisons test: *w/w* vs. *Itm2b-*KO P=0.6266, not significant. ELISA 2 sTrem2 F (2, 42) = 82.69, P<0.0001; post-hoc Tukey’s multiple comparisons test: *w/w* vs. *Itm2b-KO* P<0.0001. ELISA 2 Trem2+sTrem2 in S100 F (2, 6) = 21.06, P<0.0001; post-hoc Tukey’s multiple comparisons test: *w/w* vs. *Itm2b-*KO P=0.0703, not significant. All *w/w* vs. *Trem2-KO* and *Itm2b-KO* vs. *Trem2-KO* comparisons have P<0.0001. (c) Western blot analysis of P100 fractions from a representative *w/w, Trem2-KO* and *Itm2b-KO* P100 sample with αTrem2-CT and an αBri2 antibody. (d) Detection and quantification of Trem2-CTF in the P100 fraction by Western blot analysis and with Image Lab software. GAPDH was used as a loading control. Data were analyzed by two–way ANOVA followed by post-hoc Sidak’s multiple comparisons test when ANOVA showed significant differences. Trem2-CTF: sex factor, F (1, 28) = 1.584, P=0.2186; genotype factor, F (1, 28) = 33.25, P<0.0001; sex/genotype interaction, F (1, 28) = 2.682, P=0.1127; post-hoc Sidak’s multiple comparisons test: females, *w/w* vs. *Itm2b-KO* P<0.0001; males, *w/w* vs. *Itm2b-KO* P=0.0072. GAPDH did not show any significant differences. (e) ELISA measurements of endogenous Aβ40 and Aβ42 in brain homogenates of *w/w, Trem2-KO* and *Itm2b-KO* animals. Data were analyzed by ordinary one-way ANOVA followed by post-hoc Tukey’s multiple comparisons test when ANOVA showed significant differences. Aβ40: F (2, 36) =36.34, P<0.0001; post-hoc Tukey’s multiple comparisons test: *w/w* vs. *Itm2b-KO,* P=0.0001; *w/w* vs. *Trem2-KO,* P=0.0001; *Itm2b-KO* vs. *Trem2-KO,* P<0.0001. Aβ42: F (2, 36) =26.40, P<0.0001; post-hoc Tukey’s multiple comparisons test: *w/w* vs. *Itm2b-KO,* P=0.0003; *w/w* vs. *Trem2-KO,* P=0.0021; *Itm2b-KO* vs. *Trem2-KO,* P<0.0001. All data are shown as means +/-SEM: P=0.0005. *=P<0.05, **=P<0.01, ***=P<0.001, ****=P<0.0001.

Although Trem2-CTF cannot be detected by ELISA 1 and 2 (Figure 5a), we were able to detect it in the P100 fraction by WB analysis (Figure 5c) as it is not glycosylated. Quantification of Trem2-CTF levels in the P100 brain fractions showed that both female and male *Itm2b-KO* brains contained significantly higher steady-state levels of Trem2-CTF compared to control WT animals (Figure 5d). In summary, the absence of BRI2 in *Itm2b-KO* mice leads to increased levels of sTrem2 and Trem2-CTF, which suggests a reduction in Trem2 processing by α-secretase. However, the levels of Trem2 itself remain unchanged in *Itm2b-KO* brains, which suggests that compensatory mechanisms are in play *in vivo* to maintain Trem2 levels.

Consistent with previous reports that BRI2 reduces Aβ production by inhibiting APP cleavage^14, 15^, Aβ40 and Aβ42 levels were increased in *Itm2b-KO* mice compared to controls (Figure 5e). These findings suggest that Bri2 can inhibit both Trem2 and APP cleavage, and loss of BRI2 function leads to increased processing of both proteins. *Trem2-KO* mice, on the other hand, had significantly lower CNS Aβ40 and Aβ42 levels (Figure 5e). This finding may appear contradictory to the notion that TREM2 mediates Aβ clearance^39, 40^.

In summary, loss of Bri2 expression *in vivo* causes an increase of Trem2-CTF and sTrem2, the two products of α-secretase-processing of Trem2, which suggests an increase in Trem2 processing by α-secretase. This data, together with the evidence that BRI2 overexpression in cells lines causes a decrease in Trem2-CTF and sTrem2 levels (Figure 4), suggests that, physiologically, BRI2 dampens α-secretase-mediated processing of TREM2.

### Microglia-specific reduction of *Itm2b* increases CNS sTrem2 levels

The data in transfected cells suggest that the effect of BRI2 on Trem2 is cell-autonomous. If this were the case, reducing Bri2 expression *in vivo* only in microglia should cause an increase in sTrem2 levels. To determine the cell-autonomous or non-cell autonomous nature of Bri2’s effect on sTrem2 levels *in vivo*, we took advantage of *Cx3cr1^CreER/wt^*^41^ and *Itm2b*-Floxed (*Itm2b^f/f^*)^7, 14^ mice. The *Cx3cr1^CreER/wt^* animals contain a modified version of the chemokine (C-X3-C) receptor 1 (Cx3cr1) gene, with an inserted CreERT2 sequence followed by an internal ribosome entry site and an enhanced yellow fluorescent protein (EYFP). This results in the expression of Cre-ERT2 and EYFP only in microglia in the brain. The Cre-ERT2 fusion protein requires the presence of tamoxifen to translocate from the cytosol to the nucleus, where it can mediate loxP-loxP recombination. In *Itm2b^f/f^* mice, exon 3 of the Itm2b gene is flanked by two loxP sites. Therefore, in *Itm2b^f/f^:Cx3cr1^CreER/wt^* animals, tamoxifen administration should induce Cre-ERT2-mediated conversion of *Itm2b^f^* alleles into *Itm2b-KO* alleles, leading to suppression of *Itm2b* expression specifically in microglia but not in other brain cell types.

To verify that Cre-ERT2 and EYFP are only expressed in microglia, we prepared cell suspensions from brain tissue isolated from ∼380 days old *Cx3cr1^CreER/wt^* an*d Cx3cr1^wt/wt^* animals after intracardiac catheterization and perfusion. Cells were stained with the microglia-specific anti-CD11b-APC-conjugated antibody and analyzed by Fluorescence-activated cell sorting (FACS). The vast majority of *Itm2b^wt/wt^:Cx3cr1^CreER/wt^* microglia (CD11b^+^) were EYFP^+^ and >99% of EYFP^+^ cells were CD11b^+^ (Figure 6a), confirming that the Cre-ERT2 and EYFP expression is indeed restricted to microglia in the brain.

**Figure 6.**
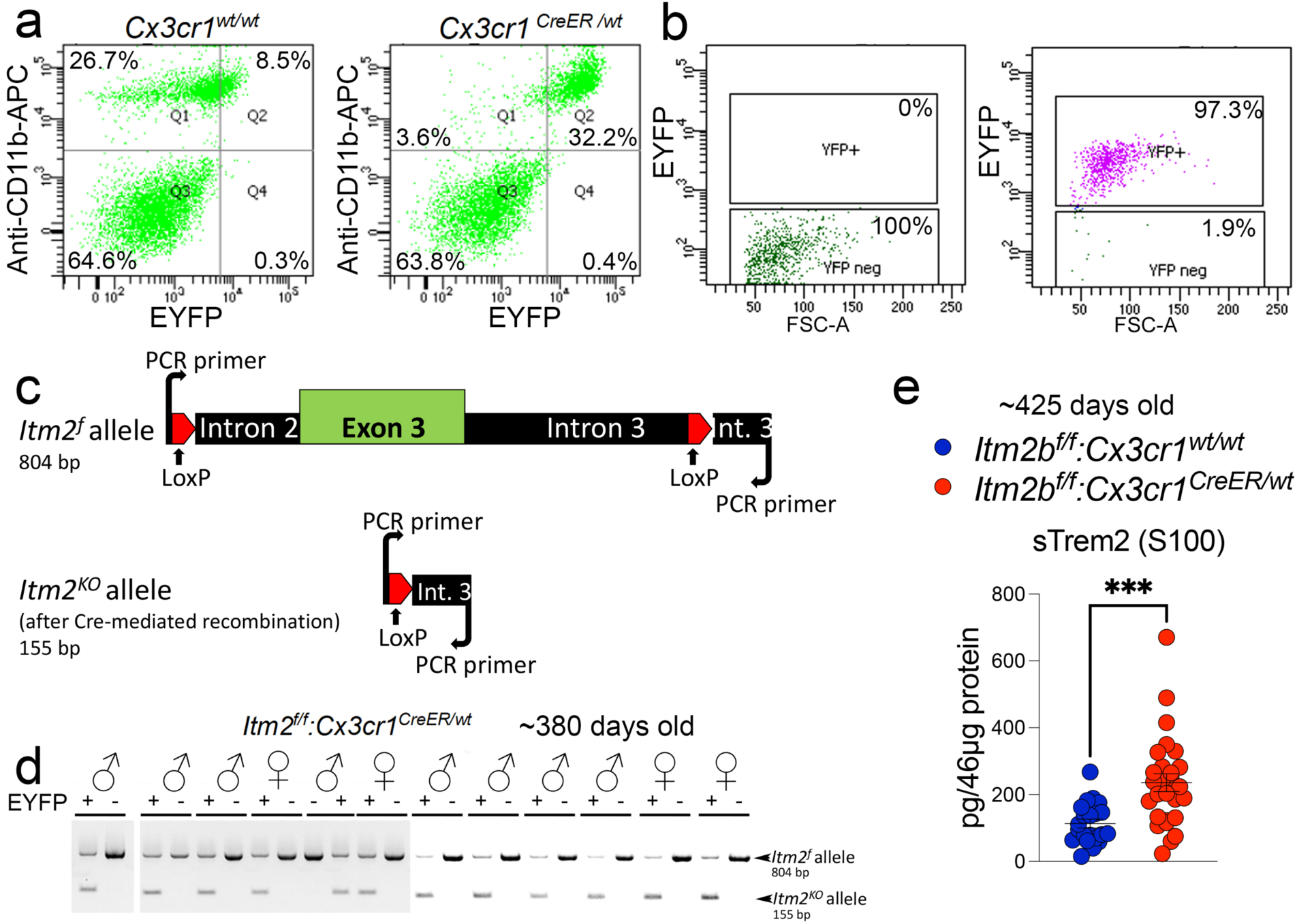
Bri2 increases CNS levels of sTrem2 via a cell-autonomous mechanism. (a) CD11b staining and FACS analysis of brain cells isolated from *Cx3cr1^CreER/wt^* an*d Cx3cr1^wt/wt^* animals. (b) FACS analysis of sorted EYFP^+^ (microglia) and EYFP^-^ (non-microglia) brain cell populations from *Itm2b^f/f^:Cx3cr1^CreER/wt^* animals. (c) Schematic representation of the PCR test used to identify the *Itm2b^f^* and *Itm2b^KO^* alleles. (d) PCR analysis of genomic DNA isolated from EYFP^+^ and EYFP^-^ cells sorted from *Itm2b^f/f^:Cx3cr1^CreER/wt^* brains. (e) ELISA 2 was used to measure sTrem2 levels in *Itm2b^f/f^:Cx3cr1^CreER/wt^* (females=15, males=12) and *Itm2b^f/f^:Cx3cr1^wt/wt^*(females=10, males=11) littermates. Data were analyzed by two-tailed unpaired t test: P=0.0004. All data are shown as means +/-SEM: ***=P<0.001.

CreERT2 mouse lines exhibit some degree of leakiness, which causes tamoxifen independent Cre activity^42^. To test whether *Itm2b^f/f^:Cx3cr1^CreER/wt^* animals showed tamoxifen independent Cre recombinase activity, we perfused ∼380 days old *Itm2b^f/f^:Cx3cr1^CreER/wt^* animals and prepared cell suspensions from brain tissue. After sorting the cells into EYFP^+^ (microglia) and EYFP^-^(non-microglia) cell populations (Figure 6b), the genomic DNA was isolated and analyzed by PCR tests to amplify the *Itm2b^f^* and the recombined *Itm2b^KO^* alleles (Figure 6c). All microglia samples (EYFP^+^) analyzed showed the presence of both the *Itm2b^f^* and *Itm2b^KO^*alleles (Figure 6d), indicating tamoxifen independent Cre recombinase activity in microglia. Non-microglia samples (EYFP^-^) showed only the *Itm2b^f^* allele (Figure 6d), indicating that Cre-ERT2 expression and partial *Itm2b* inactivation is restricted to microglia. Since the PCR method used was not quantitative, the percentage of *Itm2b^f^* alleles that had undergone recombination-conversion to *Itm2b^KO^* alleles could not be determined.

As we observed a microglia-specific partial loss of Bri2 function in *Itm2b^f/f^:Cx3cr1^CreER/wt^* mice without tamoxifen treatment, we measured sTrem2 levels in ∼425 days old *Itm2b^f/f^:Cx3cr1^CreER/wt^*and *Itm2b^f/f^:Cx3cr1^wt/wt^* littermates, without tamoxifen treatment. We found that sTrem2 levels were significantly increased in *Itm2b^f/f^:Cx3cr1^CreER/wt^* as compared to *Itm2b^f/f^:Cx3cr1^wt/wt^*littermates (Figure 6e). The increase in sTrem2 levels in *Itm2b^f/f^:Cx3cr1^CreER/wt^* mice without tamoxifen treatment indicates that the partial loss of Bri2 function in microglia alone is sufficient to increase sTrem2 levels. This data supports the idea that Bri2 inhibits α-secretase processing of Trem2 through a cell-autonomous mechanism, possibly mediated by the Bri2/Trem2 interaction. Nevertheless, it cannot be ruled out that Bri2 may also impact sTrem2 levels through a non-cell-autonomous mechanism.

## DISCUSSION

The high expression of *ITM2B*, an AD-related familial dementia gene^1–4^, in both mouse and human microglia lead to the investigation of potential functional interactions between BRI2 and TREM2 in microglia, given the emerging importance of microglia and the microglia-specific gene *TREM2* in Alzheimer’s disease pathogenesis^26, 27^. To investigate these potential interactions, we employed a multi-faceted approach, including scRNAseq analysis, biochemical and molecular assays in transfected cells and mouse models. Our findings strongly support the hypothesis that BRI2 interacts with TREM2 and modulates the processing of TREM2 in microglia in a cell-autonomous fashion.

Using scRNAseq analysis, we identified a microglial subpopulation (cluster 3) that is almost exclusively represented in *Itm2b-KO* mice. The increase in cluster 3 in *Itm2b-KO* mice, but not *Itm2b/Trem2-dKO* mice, suggests that *Itm2b* may act epistatically to *Trem2* to regulate cluster 3 expression and that Bri2 inhibits Trem2’s function: this inhibition is relieved in the absence of Bri2, leading to the expansion of clusters 3. Thus, we named this cluster the *Itm2b*-*Trem2* dependent cluster (I/T-D).

By which mechanism can Bri2 inhibit Trem2 function? BRI2 interacts with a region of APP containing the secretases cleavage sites and inhibits APP processing^14, 20^. As Trem2 is also processed by the α-secretase^28^, we hypothesized that BRI2 may inhibit Trem2 processing in a manner similar to APP. This hypothesis was tested in transiently transfected cells, and the results showed that BRI2 binds Trem2 and inhibits its processing by α-secretase, as shown by an increase of the α-secretase substrates Trem2 and a simultaneous decrease in the α-secretase products Trem2-CTF and sTrem2, and that BRI2’s inhibition of α-secretase processing of Trem2 is correlated with its binding to Trem2.

Next, we tested the physiological relevance of these findings *in vivo*, and found that mice lacking Bri2 expression had increased levels of sTrem2 and Trem2-CTF in the brain. This suggests that Bri2 plays a role in regulating Trem2 processing in vivo.

The transient transfection experiments indicate that this effect of BRI2 is cell-autonomous and depends on the BRI2-Trem2 interaction. To test this hypothesis *in vivo*, we measured sTrem2 levels in mice with reduced Bri2 expression solely in microglia and found that these mice had increased brain levels of sTrem2 compared to control mice. This suggests that the interaction between BRI2 and Trem2 is indeed cell-autonomous and plays a role in regulating Trem2 processing specifically in microglia. Altogether, the data presented in this study strongly support the hypothesis that the expansion of the I/T-D cluster in *Itm2b-KO* mice is a direct consequence of increased Trem2 processing, highlighting the physiological relevance of the Bri2-Trem2 functional interaction.

The processing of Trem2 and its functional consequences remain poorly understood, but studies suggest that Trem2 cleavage may play a crucial role in regulating the activity of microglia in the brain and in AD pathogenesis^38^. Indeed, elevated levels of sTREM2 in the cerebrospinal fluid and CNS of AD patients suggest that the processing of TREM2 by α-secretase is involved in the pathology of AD^30–32, 37^. Interestingly, FBD and FDD have a prominent neuroinflammatory component, and our findings suggest that dysregulation of the functional interaction between BRI2 and TREM2 may contribute to the neuroinflammation observed in these diseases.

The findings of this study highlight the potential of BRI2 as a therapeutic target for AD and related dementias. BRI2’s ability to regulate the processing of both APP and TREM2 suggests that it plays a multifaceted role in the pathogenesis of these diseases. Given the current lack of effective treatments for AD and related dementias, the identification of new therapeutic targets is critical. The potential of BRI2 as a therapeutic target deserves further investigation, and the development of drugs that modulate BRI2 function may provide a novel avenue for the treatment of these devastating diseases.

## Materials and methods

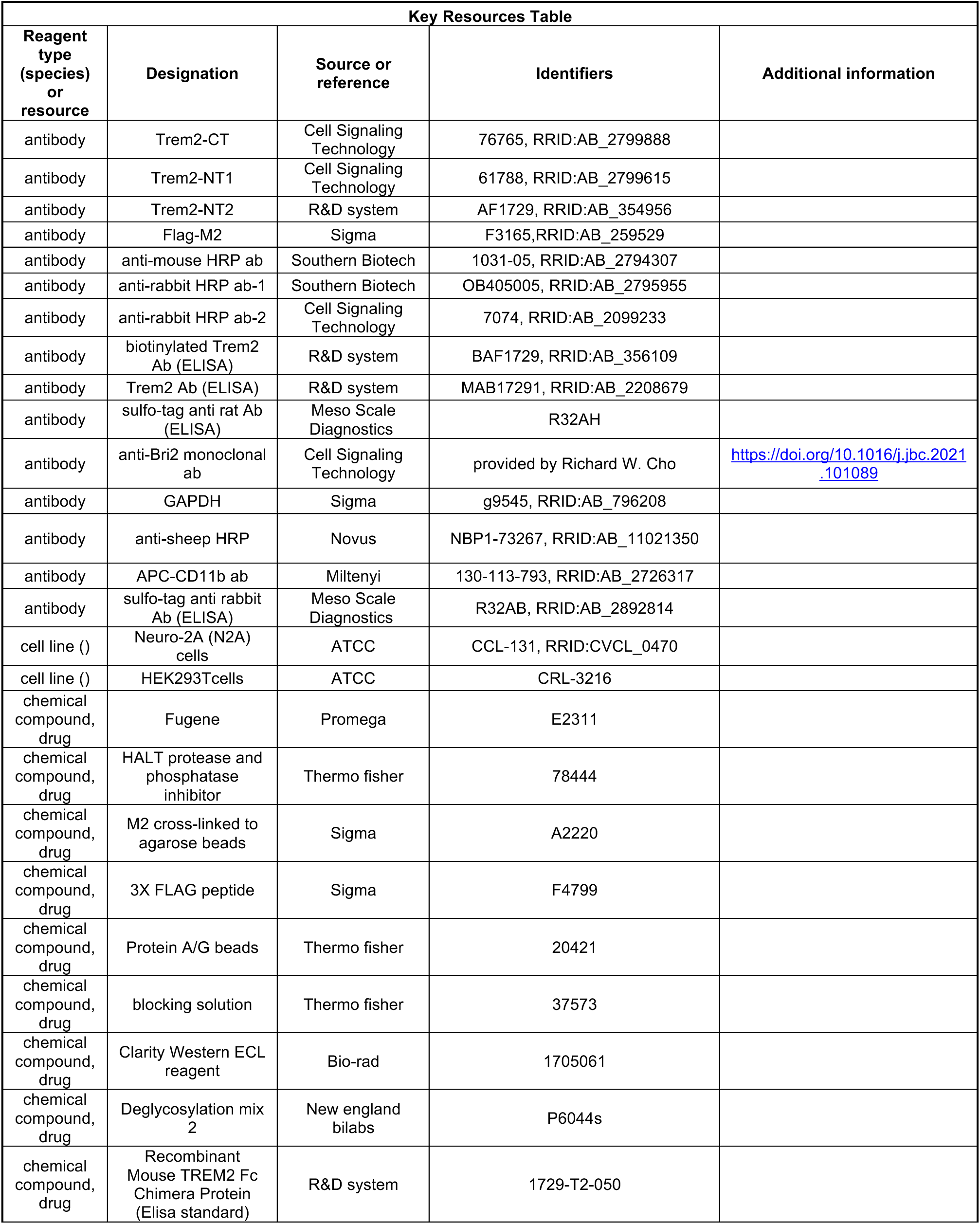
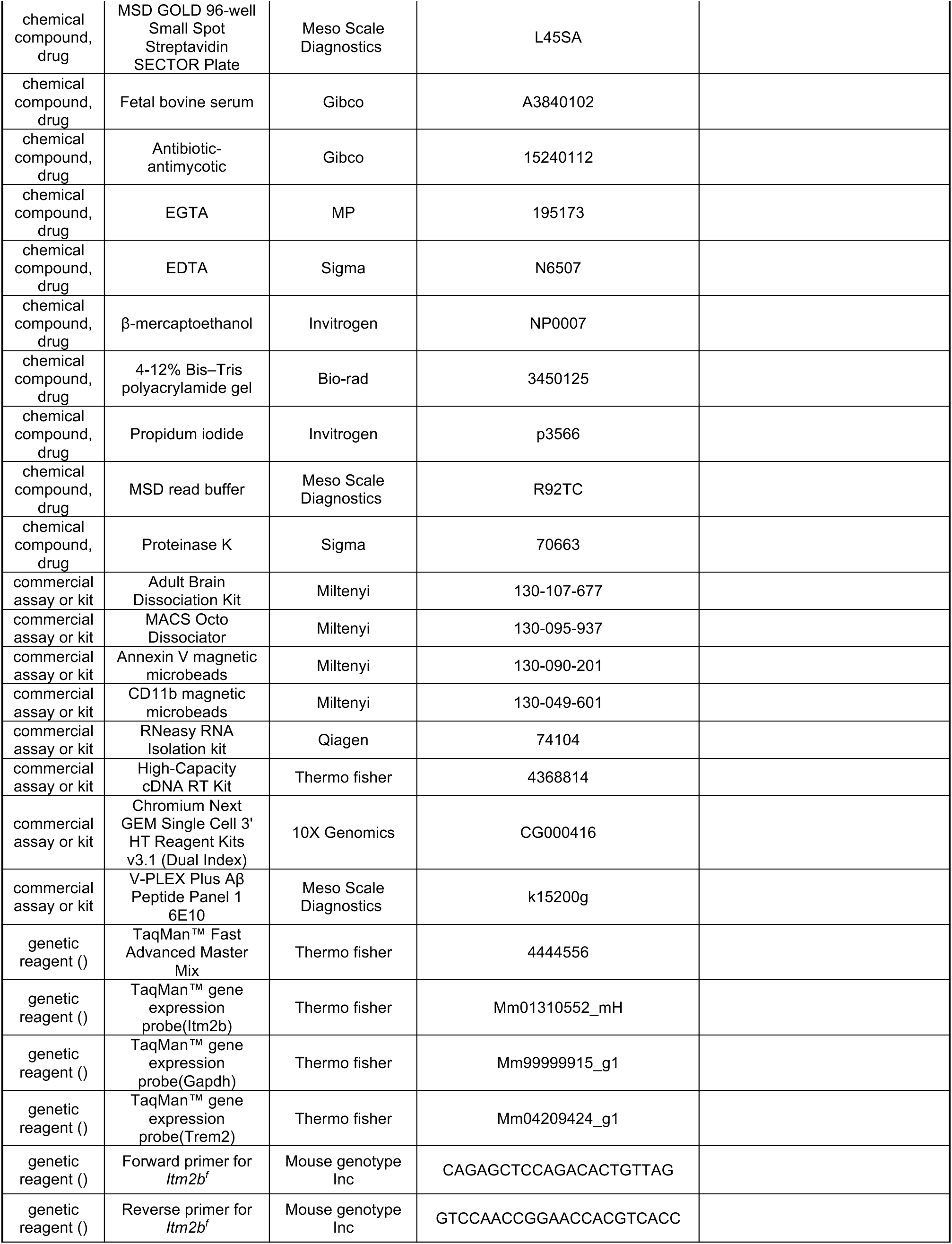
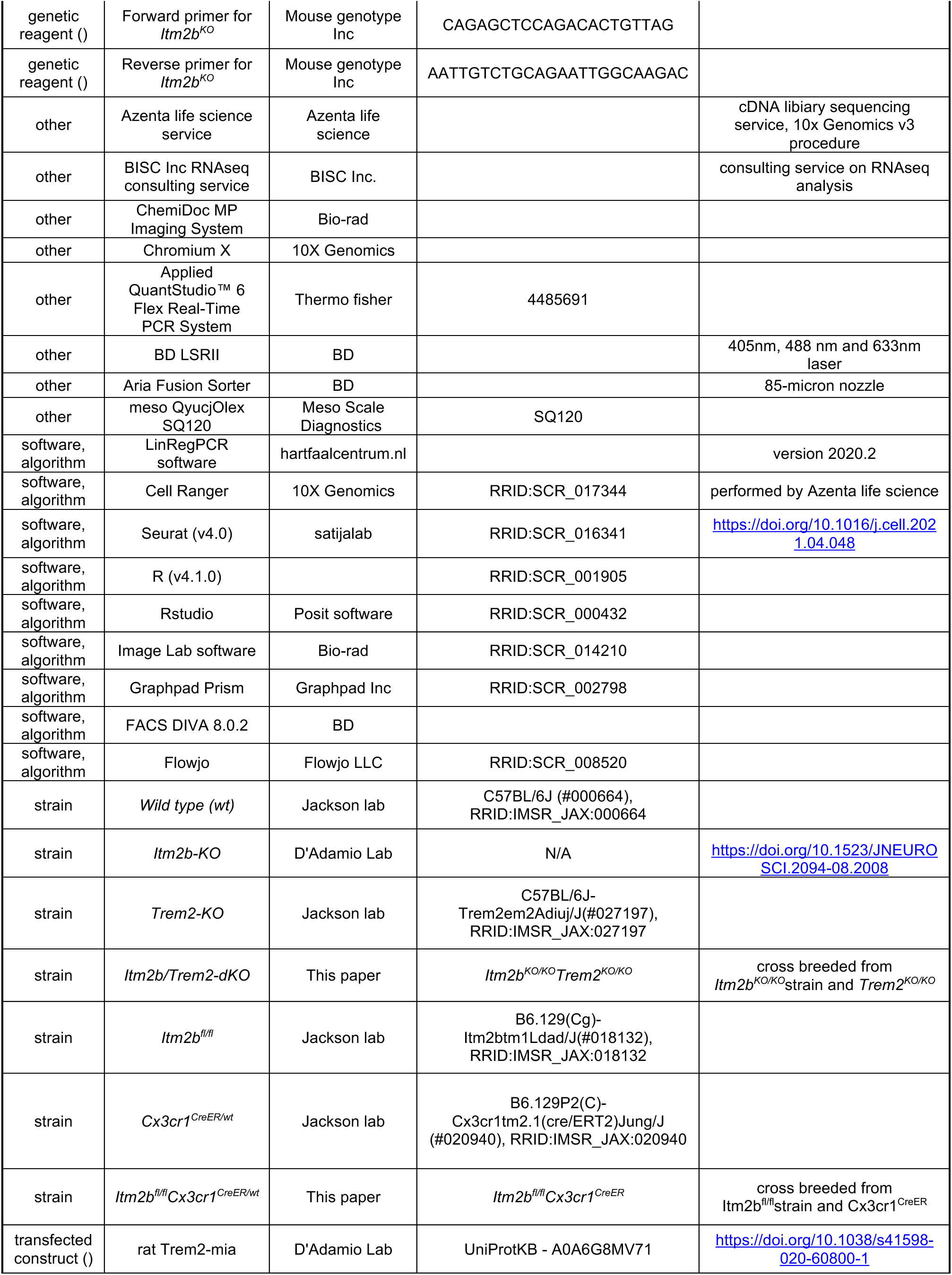
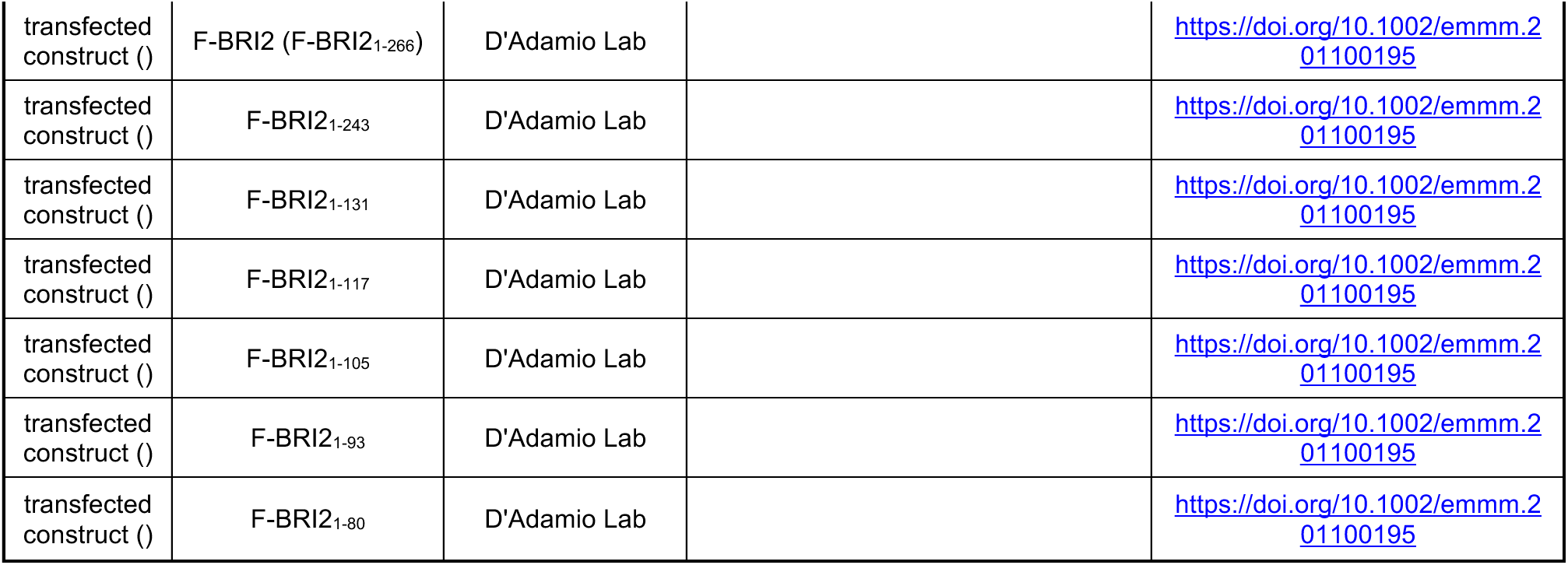

### Mice and ethics statement

All experiments were done according to policies on the care and use of laboratory animals of the Ethical Guidelines for Treatment of Laboratory Animals of the NIH. Relevant protocols were approved by the Rutgers Institutional Animal Care and Use Committee (IACUC) (Protocol #201702513). All efforts were made to minimize animal suffering and reduce the number of mice used. *Cx3cr1^CreER/wt^* and *Trem2-KO* mice were purchased from The Jackson laboratory (Stock No. 020940 and 027197, respectively). Wild type, *Itm2b-KO* and *Itm2b-floxed* animals were generated by our laboratory.

### Microglia isolation

Mouse brains were extracted from 15-month-old mice (3 females and 3 males) after intracardiac PBS perfusion to remove blood from brain blood vessels. Brains were enzymatically and mechanically dissociated into a cell suspension using the Adult Brain Dissociation Kit (Miltenyi 130-107-677) and gentle MACS Octo Dissociator (Miltenyi 130-095-937). Apoptotic cells were removed using Annexin V magnetic microbeads (Miltenyi 130-090-201) and microglia were isolated using CD11b magnetic microbeads (Miltenyi 130-049-601) according to the manufacturer’s instructions.

### Quantitative RT-PCR

Bound cells (microglia) and unbound cells (flow through, non-microglia fraction) were analyzed by quantitative RT-PCR. Total RNA was extracted from isolated cells (microglia or non-microglia cells) with RNeasy RNA Isolation kit (Qiagen 74104) and used to generate cDNA with a High-Capacity cDNA Reverse Transcription Kit (Thermo 4368814). 10 ng of cDNA, TaqMan™ Fast Advanced Master Mix (Thermo 4444556), and the appropriate TaqMan probes were used in the real time polymerase chain reaction. Samples were analyzed on an Applied QuantStudio™ 6 Flex Real-Time PCR System, and relative RNA amounts were quantified using LinRegPCR software (hartfaalcentrum.nl). The probes Mm01310552_mH (exon junction 1–2) and Mm04209424_g1 (exon junction 3-4) were used to detect mouse *Itm2b* and *Trem2*. Expression of *Itm2b* and *Trem2* were normalized to *Gapdh* levels, as detected with Mm99999915_g1 (exon junction 2–3) probe.

### ScRNAseq data generation and analysis

Microglia for scRNAseq analysis were prepared from 9–12-month-old mice in two independent experiments. Experiment-1 (Data 1) included 1 male and 1 female WT (controls), 1 male and 1 female *Itm2b-KO*, 1 male and 1 female *Trem2-KO*, 1 male and 1 female *Itm2b/Trem2-dKO*. Experiment-2 (Data 2) included 1 female WT, 1 male and 1 female *Itm2b-KO*. Microglia were isolated as described above.

Purified microglia were used to generate Gel-bead-in-emulsion (GEMs) containing single cells using Chromium X; cDNA libraries were generated following Chromium Next GEM Single Cell 3’ HT Reagent Kits v3.1 (Dual Index) instruction (10x Genomics). Sequencing was performed by Azenta life science following standard 10x Genomics v3 procedure. Raw data were pre-filtered through Cell Ranger Software. The pre-filter scRNAseq data were further analyzed using the most updated Seurat (v4.0) package in R (v4.1.0)^43^ with assistance from BISC Inc. Sequencing data for experiments-1 and -2 were assembled into two Seurat objects (called Data-1 and Data-2, respectively), which were constructed using the CreateSeuratObject function. Clusters requirements were a minimum of 3 cells and 1000 features. For quality control, cells with more than 5% mitochondrial content and/or more than 45% ribosomal content were removed. Cells with low UMI and gene number per cell were filtered out. Cutoffs for UMI and gene number were empirically determined based on histograms showing cell density as a function of UMI per gene counts. Cutoffs of UMI greater than 300 and less than 50000, and genes greater than 1000 and less than 6000 were applied to eliminate any potential subclusters formed solely due to the cells having insufficient features present to be accurately categorized while also eliminating potential doublets. We determined the dimensionality of each object to retain for downstream analysis in a heuristic manner, using an Elbow plot to establish the number of principal components (PCs) necessary to capture 95% of the variance in gene expression (Supplementary Figure 4). We next constructed a K-nearest neighbors (KNN) graph based on the Euclidean distance in PCA space, refining the edge weights between any two cells based on the shared overlap in their local neighborhoods (a measure known as the Jaccard similarity) with the FindNeighbors() function^43^. To cluster the cells, we then applied a modularity optimization technique (using the Louvain algorithm) implemented in the FindClusters() function (using a resolution of 0.4). Datasets were visualized in two dimensions using a uniform manifold approximation and projection (UMAP) dimensional reduction (https://doi.org/10.21105/joss.00861).

To specifically select microglia for further analysis, we performed cell type annotation using a single cell dataset published by Van Hove^35^ as a reference. First, the Van Hove reference dataset was re-normalized using the SCTransform (v2) to be compatible with Data1 and Data2, before finding mutual nearest neighbor gene “anchors” between the reference and the Data1 and Data2 objects respectively using the FindTransferAnchors function. Next, we used this set of anchors to predict the identities of the Data1/Data2 cells. Cells predicted to be of the type “microglia” with > 95% confidence were retained for further analysis.

Next, we evaluated the effects of *Trem2* and/or *Itm2b* deletion on microglia’s gene expression. However, as the samples were sequenced in two different experiments, the scRNA dataset integration functionality of the Seurat package was used to perform the joint analysis as previously described. This resulted in an object containing information on 297,215 cells. The top 3,000 most highly variable genes of each dataset were detected by the SelectIntegrationFeatures function to use as feature anchors in the PrepSCTIntegration function. The FindIntegrationAnchors function was then run on the SCT(v2) select sample datasets indicated above from Data1 and Data2 were integrated using the first 20 principal components into Object1.

Principal component analysis (PCA) was reperformed with PCA function before carrying out further sequencing batch correction normalization with Harmony^44^. Clustering of the integrated data was then executed using the top 30 harmonized PCA components to find neighbors and an SNN clustering resolution of 0.4. This resolution appeared to produce the most informative microglial clustering, but the microglia subtype investigation was generally robust to various choices of this hyperparameter.

### Kegg pathway analysis

Cluster 3 pathway analysis was performed using DEenrichRPlot function referenced with KEGG_2019_Mouse database.

### Cell culture, plasmids, transfection, immunoprecipitation and Western blots

**analysis.** Neuro-2A (N2A) cells (ATCC CCL-131) and HEK293Tcells (ATCC CRL-3216) were maintained in Eagle’s Minimum Essential Medium (EMEM) (Gibco 11095-098) supplemented with 10% fetal bovine serum (Gibco A3840102) and Antibiotic-Antimycotic (Gibco 15240112). Plasmids used where described previously in the included references. Rat Trem2 (the Trem2-Miα isoform, UniProtKB - A0A6G8MV71)^38^; F-BRI2 (A.K.A. F-BRI2_1-266_), F-BRI2_1-243_, F-BRI2_1-131_, F-BRI2_1-117_, F-BRI2_1-105_, F-BRI2_1-93_ and F-BRI2_1-80_ ^12, 45^. Both HEK293T cells and N2A cells were transfected with indicated plasmids *via* Fugene (Promega, E2311) as previously described^46, 47^.

Transfected cell lysate was lysed in immunoprecipitation buffer (50 mm Tris, 150 mm NaCl, 1 mm EGTA (MP 195173), and 1 mm EDTA (pH 8.0) (Sigma E4378) with 0.5% NP40 (Sigma N6507) and HALT protease/phosphatase inhibitor (Thermo fisher 78444) solubilized for 30 min at 4 °C with rotating and spun at 20,000 × g for 10 min. Solubilized cell lysate was used as input for immunoprecipitation. FLAG-BRI2 proteins were immunoprecipitated with anti-FLAG mouse monoclonal antibody M2 cross-linked to agarose beads (Sigma A2220); immunoprecipitated proteins were eluted with 3X FLAG peptide (Sigma F4799). Trem2 was immunoprecipitated with either a Rabbit monoclonal antibody raised against the carboxy-terminus of mouse Trem2 (Cell Signaling Technology, 76765,) referred to as αTrem2-CT, a Rabbit monoclonal antibody raised against the amino-terminus of mouse Trem2 (CST 61788) αTrem2-NT2, or the sheep polyclonal IgG raised against the amino-terminus of mouse Trem2 anti (R&D AF1729) αTrem2-NT1. Immunocomplexes were isolated with protein A/G beads (Thermo, 20421) and eluted 1× LDS sample buffer with 10% β-mercaptoethanol (Invitrogen; NP0007) at 55 °C. Input (total lysates, T.L.) and immunoprecipitation (I.P.) eluates were analyzed by Western blot.

For Western blot analyses, total lysates proteins were diluted with PBS and LDS sample buffer—10% β-mercaptoethanol (Invitrogen; NP0007), separated on a 4 to 12% Bis–Tris polyacrylamide gel (Bio-Rad; 3450125), and transferred onto nitrocellulose at 25 V for 7 min using the Trans-blot Turbo system (Bio-Rad). Blotting efficiency was visualized by red Ponceau staining on membranes. Membranes were blocked in 5% milk (Bio-Rad; 1706404) for 45 min and washed in PBS/0.05% Tween-20. Primary antibodies were applied in blocking solution (Thermo; 37573). The following antibodies were used: anti-Flag M2 (Sigma F3165), anti-Trem2-CT, anti-Trem2-NT1, anti-Bri2 monoclonal antibody (provided by Richard W. Cho Cell Signaling Technology), anti-GAPDH (Sigma g9545). Secondary antibodies [either anti-mouse (Southern Biotech; 1031-05), anti-sheep (Novus NBP1-73267) or a 1:1 mix of anti-rabbit (Southern Biotech; OB405005) and anti-rabbit (Cell Signaling; 7074)] were diluted 1:1000 in 5% milk and used against either mouse or rabbit primary antibodies for 1 h at room temperature, with shaking. Membranes were washed with PBS/Tween-20 to 0.05% (three times, 10 min each time), developed with Clarity Western ECL reagent (Bio-rad 1705061) and visualized on a ChemiDoc MP Imaging System (Bio-Rad). Signal intensity was quantified with Image Lab software (Bio-Rad). Data were analyzed using Prism software (GraphPad Software, Inc) and represented as mean ± SEM.

### Deglycosylation prior to Western blot analysis

Cell lysate or total microglia were solubilized with 1% NP-40 for 30 min rotating, spun at 20,000 g and the supernatant was used as input for deglycosylation reactions, according to the manufacturer’s specifications (NEB P6044S). Conditioned media media were deglycosylated directly, with no prior solubilization step.

### Flow cytometry

Cells were dissociated from adult mouse brains as described above and prepared in FACS buffer (PBS w/w 2% BSA + 1 mM EDTA). The cells were stained with APC-CD11b antibody (Miltenyi 130-113-793) for 30 min with three subsequent washes. Propidium iodide (1%, Invitrogen p3566) was added to eliminate dead cells from analysis. Cells were acquired using a LSRII (BD Bioscience) and DIVA 8.0.2 software, and data were analyzed using FlowJo software.

### Cell sorting

Cells were dissociated from adult mouse brains as described above. Propidium iodide (1%, Invitrogen p3566) was added to a single-cell suspension before sorting EYFP^+^ and EYFP^-^ cells using a FACS Aria Fusion Sorter (BD Bioscience). To verify sorting efficiency, sorted cells were acquired using a LSRII (BD Bioscience) and DIVA 8.0.2 software, and data were analyzed using FlowJo software.

### ELISAs

For analysis of human Aβ peptides, the brain lysates were diluted at 4 µg/µl. Aβ38, Aβ40, and Aβ42 were measured using a V-PLEX Plus Aβ Peptide Panel 1 6E10 (Meso Scale Discovery, K15200G). The plates were read on a MESO QuickPlex SQ 120. The ELISAs for analysis of full length Trem2 and sTrem2 were modified from Kleinberger’s protocol^48^. Briefly, a streptavidin-coated plate (Meso Scale Discovery, L15SA) was blocked with 3% BSA/PBST (0.05% Tween-20) overnight at 4 °C. The plates were incubated with 0.25 μg/ml biotinylated sheep anti-mouse TREM2 antibody (R&D Systems, BAF1729) for 1 hr at RT with shaking. After washing four times with PBST, the samples and standards (Trem2, R&D Systems, 1729-T2-050) were incubated for 2 hrs at RT with shaking. The plates were washed three times with PBST and incubated with either 1 μg/ml of rabbit monoclonal anti-mouse TREM2 (Cell Signaling Technology, 76765) for full length Trem2 ELISA (ELISA 1), or with 1 μg/ml rat monoclonal anti-mouse/human TREM2 (R&D Systems, MAB17291) for sTrem2 ELISA (ELISA 2) for 1 hr at RT. After four washes in PBST, 0.5 µg/ml SULFO-TAG labeled antibody (Meso Scale Discovery, anti-rabbit R32AB for ELISA 1; anti-rat R32AH for ELISA 2) was added and incubated for 1 hr with shaking. The plates were washed three times in PBST, developed in Meso Scale Discovery read buffer (Meso Scale Discovery, R92TC), and read on a MESO QuickPlex SQ 120.

### Genomic DNA isolation and PCR analysis

FACS sorted EYFP^+^ cells were incubated in 300μl of lysis buffer (100mM Tris, 5mM EDTA, 0.2% SDS, 200mM NaCl, PH 8.0) with 60 μg/ml of protease K (Sigma 70663) at 55^0^ C for 2 hrs. 100 μl of protein precipitation solution (7.5M Ammonium Acetate) was added and vortex for 30s before centrifugation at 15000xg for 5 min. The supernatant was transferred into a new Eppendorf tube with 300 μl Isopropanol and mixed by inverting 30 times. The samples were subsequently centrifuged at 15000xg for 5 min and washed with 70% EtOH. Dried tube with genomic DNA were resuspend in 100 μl of water for PCR analysis. Primer pairs were: Forward primer-CAGAGCTCCAGACACTGTTAG, Reverse primer-GTCCAACCGGAACCACGTCACC to amplify the *Itm2b^f^* allele (804 bp PCR product); Forward primer-CAGAGCTCCAGACACTGTTAG, Reverse primer-AATTGTCTGCAGAATTGGCAAGAC to amplify the *Itm2b^KO^* allele (155 bp PCR product). The PCR was performed as follows: denaturation at 94^0^C for 3min, followed by 34 cycles of denaturation at 94^0^C for 30s, annealing at 58^0^C for 30s, extension at 72^0^C for 40s, followed by a final “filling” extension at 72^0^C for 2min. PCRs were performed by Mouse genotype Inc.

## LIST OF ABBREVIATIONS

AD: Alzheimer’s disease
FDD: Familial Danish Dementia
FBD: Familial British Dementia
APP: Amyloid-β Precursor protein
Aβ: Amyloid β-peptide
CNS: central nervous system
scRNAseq: single cell RNAseq
snRNAseq: single nuclei RNAseq
UMAP: uniform manifold approximation and projection
sTREM2: soluble TREM2 ectodomain
TREM2-CTF: C-terminal TREM2 fragment
EYFP: enhanced yellow fluorescent protein
FACS: Fluorescence-activated cell sorting
DFC: human dorso-lateral prefrontal cortex

## DECLARATIONS

### Ethics approval and consent to participate

Ethical care and use of animals in accordance with the Ethical Guidelines for Treatment of Laboratory Animals of the NIH. The procedures were described and approved by the Institutional Animal Care and Use Committee (IACUC) at Rutgers (PROTO201702513).

### Consent for publication

Not applicable.

### Availability of data and materials

The datasets and materials used and/or analyzed during the current study are available from the corresponding author on reasonable request. The scRNAseq data are being deposited at https://www.ncbi.nlm.nih.gov/geo/info/seq.html to allow public access once the data are published.

### Competing interests

The authors declare that they have no competing interests.

### Funding

National Institute on Aging (To LD: RO1AG033007 and R01AG073182)

### Authors’ contributions

T.Y. and L.D. designed the experiments, analyzed the results, performed the experiments and wrote the manuscript.

## Acknowledgements

All authors read and approved the final manuscript.

## Authors’ information

Department of Pharmacology, Physiology & Neuroscience New Jersey Medical School, Brain Health Institute, Jacqueline Krieger Klein Center in Alzheimer’s Disease and Neurodegeneration Research, Rutgers, The State University of New Jersey, 205 South Orange Ave, Newark, NJ, 07103, USA

---

**Supplementary Figure 1.**
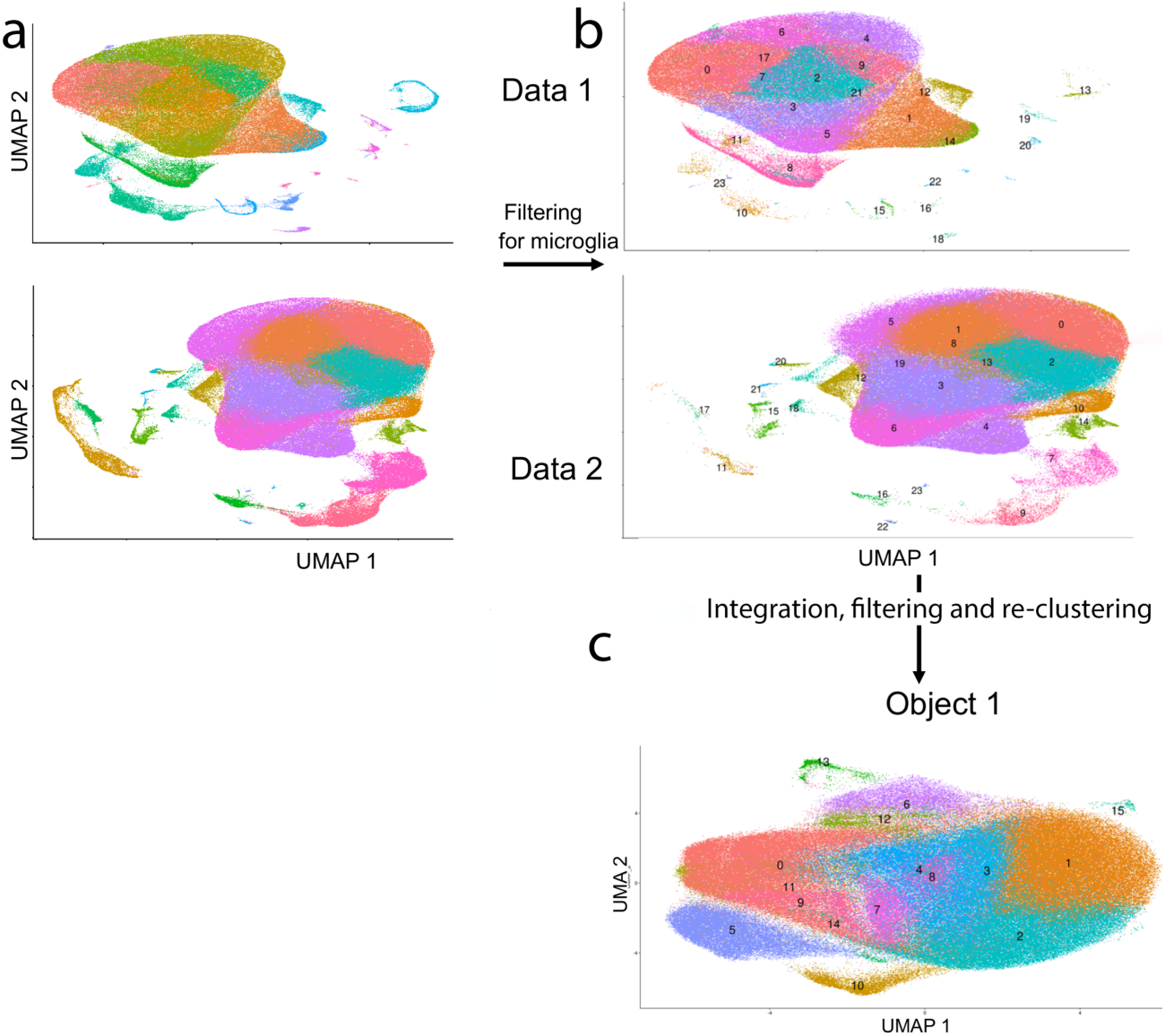
UMAP of Data 1 and Data 2 objects before (a) and after filtering for microglia (b). (c) UMAP of Object 1 after integration, filtering, and re-clustering, shows 16 microglia clusters.

**Supplementary Figure 2.**
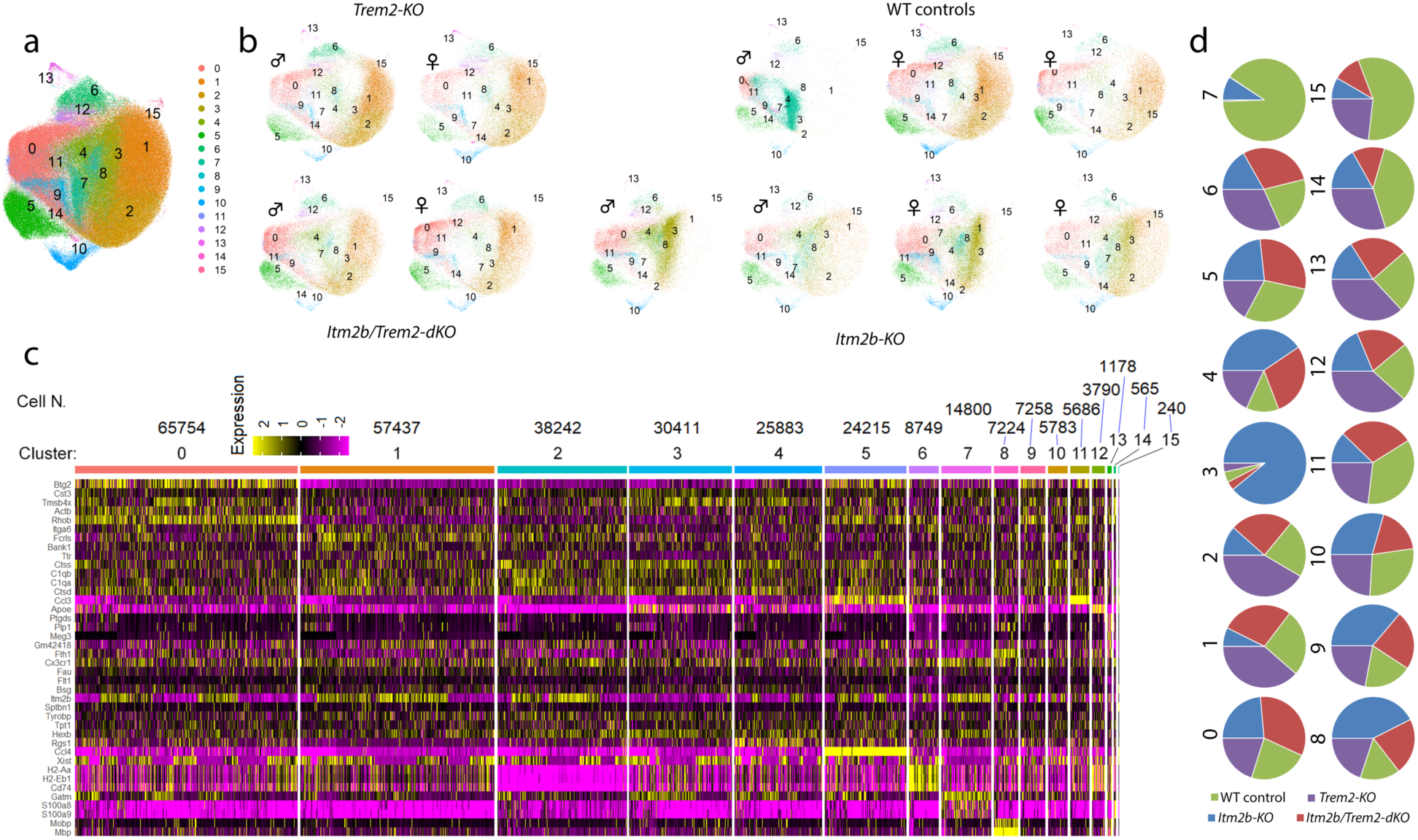
(a) UMAPs of re-clustered microglia in Object 1. (b) UMAPs split by individual samples. (c) Gene expression heatmap showing the top 5 enriched genes for each microglia cluster. The number of cells per cluster is denoted above the cluster. *Itm2b* is one of the top genes downregulated in cluster 3 because 89% of the cells in this cluster are from *Itm2b-KO* mice. (d) Proportional contribution of each genotype to each cluster. Cluster 3 was highly represented in *Itm2b-KO* mice, with 89% of microglia in this cluster originating from these mice. Conversely, Cluster 7 was preponderant in WT controls. However, ∼93% of the cells assigned to cluster 7 derived from one WT control animal (the male WT control, as depicted in UMAP plot b). Therefore, the observed expansion of cluster 7 is attributed to animal-specific factors rather than genotype-specific factors.

**Supplementary Figure 3.**
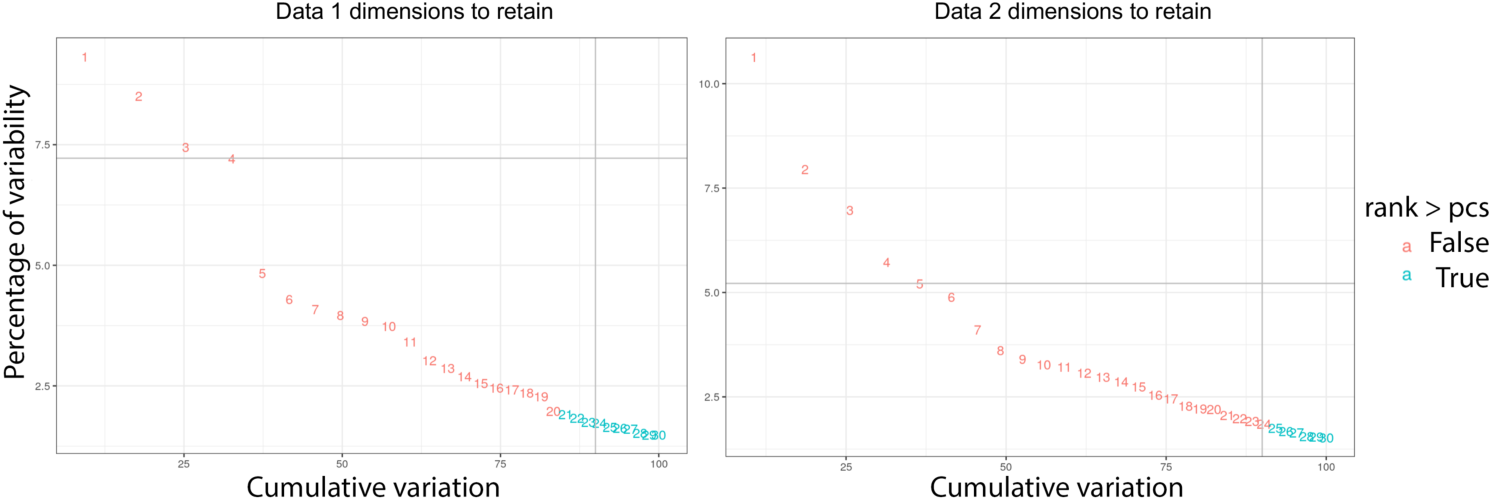
Heuristic establishment of the number of dimensions/principal components (PCs) needed to capture 90% of the variance in gene expression from the datasets.

## Notes

### Competing Interest Statement

The authors have declared no competing interest.

## References

1 Vidal, R. et al. A stop-codon mutation in the BRI gene associated with familial British dementia. Nature 399, 776–781 (1999). https://doi.org:10.1038/21637

2 Vidal, R. et al. A decamer duplication in the 3’ region of the BRI gene originates an amyloid peptide that is associated with dementia in a Danish kindred. Proc Natl Acad Sci U S A 97, 4920–4925 (2000). https://doi.org:10.1073/pnas.080076097

3 Liu, X. et al. A Novel ITM2B Mutation Associated with Familial Chinese Dementia. J Alzheimers Dis 81, 499–505 (2021). https://doi.org:10.3233/JAD-210176

4 Rhyu, J. M. et al. A Novel c.800G>C Variant of the ITM2B Gene in Familial Korean Dementia. J Alzheimers Dis (2023). https://doi.org:10.3233/JAD-230051

5 Choi, S. I., Vidal, R., Frangione, B. & Levy, E. Axonal transport of British and Danish amyloid peptides via secretory vesicles. FASEB J 18, 373–375 (2004). https://doi.org:10.1096/fj.03-0730fje

6 Garringer, H. J., Murrell, J., D’Adamio, L., Ghetti, B. & Vidal, R. Modeling familial British and Danish dementia. Brain Struct Funct 214, 235–244 (2010). https://doi.org:10.1007/s00429-009-0221-9

7 Yao, W., Yin, T., Tambini, M. D. & D’Adamio, L. The Familial dementia gene ITM2b/BRI2 facilitates glutamate transmission via both presynaptic and postsynaptic mechanisms. Sci Rep 9, 4862 (2019). https://doi.org:10.1038/s41598-019-41340-9

8 Yin, T., Yao, W., Lemenze, A. D. & D’Adamio, L. Danish and British dementia I*TM2b/BRI2* mutations reduce BRI2 protein stability and impair glutamatergic synaptic transmission. J Biol Chem (2020). https://doi.org:10.1074/jbc.RA120.015679

9 Yin, T., Yao, W., Lemenze, A. D. & D’Adamio, L. Danish and British dementia ITM2b/BRI2 mutations reduce BRI2 protein stability and impair glutamatergic synaptic transmission. J Biol Chem 296, 100054 (2021). https://doi.org:10.1074/jbc.RA120.015679

10 Tamayev, R., Matsuda, S., Fa, M., Arancio, O. & D’Adamio, L. Danish dementia mice suggest that loss of function and not the amyloid cascade causes synaptic plasticity and memory deficits. Proc Natl Acad Sci U S A 107, 20822–20827 (2010). https://doi.org:10.1073/pnas.1011689107

11 Tamayev, R. et al. Memory deficits due to familial British dementia BRI2 mutation are caused by loss of BRI2 function rather than amyloidosis. J Neurosci 30, 14915–14924 (2010). https://doi.org:10.1523/JNEUROSCI.3917-10.2010

12 Matsuda, S. et al. The familial dementia BRI2 gene binds the Alzheimer gene amyloid-beta precursor protein and inhibits amyloid-beta production. J Biol Chem 280, 28912–28916 (2005). https://doi.org:10.1074/jbc.C500217200

13 Matsuda, S., Matsuda, Y., Snapp, E. L. & D’Adamio, L. Maturation of BRI2 generates a specific inhibitor that reduces APP processing at the plasma membrane and in endocytic vesicles. Neurobiol Aging 32, 1400–1408 (2011). https://doi.org:10.1016/j.neurobiolaging.2009.08.005

14 Matsuda, S., Giliberto, L., Matsuda, Y., McGowan, E. M. & D’Adamio, L. BRI2 inhibits amyloid beta-peptide precursor protein processing by interfering with the docking of secretases to the substrate. J Neurosci 28, 8668–8676 (2008). https://doi.org:10.1523/JNEUROSCI.2094-08.2008

15 Tamayev, R., Matsuda, S., Giliberto, L., Arancio, O. & D’Adamio, L. APP heterozygosity averts memory deficit in knockin mice expressing the Danish dementia BRI2 mutant. EMBO J 30, 2501–2509 (2011). https://doi.org:10.1038/emboj.2011.161

16 Chen, G. et al. Augmentation of Bri2 molecular chaperone activity against amyloid-β reduces neurotoxicity in mouse hippocampus in vitro. Commun Biol 3, 32 (2020). https://doi.org:10.1038/s42003-020-0757-z

17 Tambaro, S., Galán-Acosta, L., Leppert, A., Presto, J. & Johansson, J. BRICHOS - an anti-amyloid chaperone: evaluation of blood-brain barrier permeability of Bri2 BRICHOS. Amyloid 24, 7–8 (2017). https://doi.org:10.1080/13506129.2016.1272451

18 Matsuda, S., Tamayev, R. & D’Adamio, L. Increased AβPP processing in familial Danish dementia patients. J Alzheimers Dis 27, 385–391 (2011). https://doi.org:10.3233/JAD-2011-110785

19 Tamayev, R. & D’Adamio, L. Inhibition of γ-secretase worsens memory deficits in a genetically congruous mouse model of Danish dementia. Mol Neurodegener 7, 19 (2012). https://doi.org:10.1186/1750-1326-7-19

20 Tamayev, R., Matsuda, S., Arancio, O. & D’Adamio, L. β-but not γ-secretase proteolysis of APP causes synaptic and memory deficits in a mouse model of dementia. EMBO Mol Med 4, 171–179 (2012). https://doi.org:10.1002/emmm.201100195

21 Tamayev, R. & D’Adamio, L. Memory deficits of British dementia knock-in mice are prevented by Aβ-precursor protein haploinsufficiency. J Neurosci 32, 5481–5485 (2012). https://doi.org:10.1523/JNEUROSCI.5193-11.2012

22 Zeisel, A. et al. Molecular Architecture of the Mouse Nervous System. Cell 174, 999–1014.e1022 (2018). https://doi.org:10.1016/j.cell.2018.06.021

23 Li, M. et al. Integrative functional genomic analysis of human brain development and neuropsychiatric risks. Science 362 (2018). https://doi.org:10.1126/science.aat7615

24 Akiyama, H. et al. Inflammation and Alzheimer’s disease. Neurobiol Aging 21, 383–421 (2000). https://doi.org:10.1016/s0197-4580(00)00124-x

25 Tarkowski, E., Andreasen, N., Tarkowski, A. & Blennow, K. Intrathecal inflammation precedes development of Alzheimer’s disease. J Neurol Neurosurg Psychiatry 74, 1200–1205 (2003). https://doi.org:10.1136/jnnp.74.9.1200

26 Schmid, C. D. et al. Heterogeneous expression of the triggering receptor expressed on myeloid cells-2 on adult murine microglia. J Neurochem 83, 1309–1320 (2002). https://doi.org:10.1046/j.1471-4159.2002.01243.x

27 Guerreiro, R. et al. TREM2 variants in Alzheimer’s disease. N Engl J Med 368, 117–127 (2013). https://doi.org:10.1056/NEJMoa1211851

28 Wunderlich, P. et al. Sequential proteolytic processing of the triggering receptor expressed on myeloid cells-2 (TREM2) protein by ectodomain shedding and γ-secretase-dependent intramembranous cleavage. J Biol Chem 288, 33027–33036 (2013). https://doi.org:10.1074/jbc.M113.517540

29 Thornton, P. et al. TREM2 shedding by cleavage at the H157-S158 bond is accelerated for the Alzheimer’s disease-associated H157Y variant. EMBO Mol Med 9, 1366–1378 (2017). https://doi.org:10.15252/emmm.201707673

30 Heslegrave, A. et al. Increased cerebrospinal fluid soluble TREM2 concentration in Alzheimer’s disease. Mol Neurodegener 11, 3 (2016). https://doi.org:10.1186/s13024-016-0071-x

31 Piccio, L. et al. Cerebrospinal fluid soluble TREM2 is higher in Alzheimer disease and associated with mutation status. Acta Neuropathol 131, 925–933 (2016). https://doi.org:10.1007/s00401-016-1533-5

32 Suárez-Calvet, M. et al. Early increase of CSF sTREM2 in Alzheimer’s disease is associated with tau related-neurodegeneration but not with amyloid-β pathology. Mol Neurodegener 14, 1 (2019). https://doi.org:10.1186/s13024-018-0301-5

33 McInnes, L., Healy, J. & Melville, J. UMAP: Uniform Manifold Approximation and Projection for dimension reduction. arXiv preprint arXiv:1802.03426 (2018).

34 Tambini, M. D. & D’Adamio, L. Trem2 Splicing and Expression are Preserved in a Human Abeta-producing, Rat Knock-in Model of Trem2-R47H Alzheimer’s Risk Variant. Sci Rep 10, 4122 (2020). https://doi.org:10.1038/s41598-020-60800-1

35 Van Hove, H. et al. A single-cell atlas of mouse brain macrophages reveals unique transcriptional identities shaped by ontogeny and tissue environment. Nat Neurosci 22, 1021–1035 (2019). https://doi.org:10.1038/s41593-019-0393-4

36 Chen, Y. & Colonna, M. Microglia in Alzheimer’s disease at single-cell level. Are there common patterns in humans and mice? J Exp Med 218 (2021). https://doi.org:10.1084/jem.20202717

37 Lichtenthaler, S. F., Tschirner, S. K. & Steiner, H. Secretases in Alzheimer’s disease: Novel insights into proteolysis of APP and TREM2. Curr Opin Neurobiol 72, 101–110 (2022). https://doi.org:10.1016/j.conb.2021.09.003

38 Tambini, M. D. & D’Adamio, L. Trem2 Splicing and Expression are Preserved in a Human Aβ-producing, Rat Knock-in Model of Trem2-R47H Alzheimer’s Risk Variant. Sci Rep 10, 4122 (2020). https://doi.org:10.1038/s41598-020-60800-1

39 Lessard, C. B. et al. High-affinity interactions and signal transduction between Abeta oligomers and TREM2. EMBO Mol Med 10 (2018). https://doi.org:10.15252/emmm.201809027

40 Yeh, F. L., Wang, Y., Tom, I., Gonzalez, L. C. & Sheng, M. TREM2 Binds to Apolipoproteins, Including APOE and CLU/APOJ, and Thereby Facilitates Uptake of Amyloid-Beta by Microglia. Neuron 91, 328–340 (2016). https://doi.org:10.1016/j.neuron.2016.06.015

41 Yona, S. et al. Fate mapping reveals origins and dynamics of monocytes and tissue macrophages under homeostasis. Immunity 38, 79–91 (2013). https://doi.org:10.1016/j.immuni.2012.12.001

42 Alvarez-Aznar, A. et al. Tamoxifen-independent recombination of reporter genes limits lineage tracing and mosaic analysis using CreER(T2) lines. Transgenic Res 29, 53–68 (2020). https://doi.org:10.1007/s11248-019-00177-8

43 Hao, Y. et al. Integrated analysis of multimodal single-cell data. Cell 184, 3573–3587 e3529 (2021). https://doi.org:10.1016/j.cell.2021.04.048

44 Korsunsky, I. et al. Fast, sensitive and accurate integration of single-cell data with Harmony. Nat Methods 16, 1289–1296 (2019). https://doi.org:10.1038/s41592-019-0619-0

45 Tamayev, R., Matsuda, S., Arancio, O. & D’Adamio, L. beta-but not gamma-secretase proteolysis of APP causes synaptic and memory deficits in a mouse model of dementia. EMBO Mol Med 4, 171–179 (2012). https://doi.org:10.1002/emmm.201100195

46 Scheinfeld, M. H. et al. Jun NH2-terminal kinase (JNK) interacting protein 1 (JIP1) binds the cytoplasmic domain of the Alzheimer’s beta-amyloid precursor protein (APP). J Biol Chem 277, 3767–3775 (2002). https://doi.org:10.1074/jbc.M108357200

47 Noviello, C., Vito, P., Lopez, P., Abdallah, M. & D’Adamio, L. Autosomal recessive hypercholesterolemia protein interacts with and regulates the cell surface level of Alzheimer’s amyloid beta precursor protein. J Biol Chem 278, 31843–31847 (2003). https://doi.org:10.1074/jbc.M304133200

48 Kleinberger, G. et al. TREM2 mutations implicated in neurodegeneration impair cell surface transport and phagocytosis. Sci Transl Med 6, 243ra286 (2014). https://doi.org:10.1126/scitranslmed.3009093

